# A novel population of long-range inhibitory neurons

**DOI:** 10.1101/554360

**Authors:** Zoé Christenson Wick, Madison R. Tetzlaff, Esther Krook-Magnuson

**Affiliations:** Graduate Program in Neuroscience – University of Minnesota; Neuroscience Department – University of Minnesota

## Abstract

The hippocampus, a brain region important for spatial navigation and episodic memory, benefits from a rich diversity of neuronal cell-types. Recent work suggests fundamental gaps in our knowledge of these basic building blocks (i.e., neuronal types) in the hippocampal circuit, despite extensive prior examination. Through the use of an intersectional genetic viral vector approach, we report a novel hippocampal neuronal population, which has not previously been characterized, and which we refer to as LINCs. LINCs are GABAergic, but, in addition to broadly targeting local CA1 cells, also have long-range axons. LINCs are thus both interneurons and projection neurons. We demonstrate that LINCs, despite being relatively few in number, can have a strong influence on both hippocampal and extrahippocampal network synchrony and function. Identification and characterization of this novel cell population advances our basic understanding of both hippocampal circuitry and neuronal diversity.

## INTRODUCTION

The hippocampus is one of the most extensively studied brain regions^1^, and in CA1 alone, over 20 types of inhibitory neurons have been previously described^2,3^. Each population of neurons plays a unique role in the circuitry^4–7^, and together, allow for the emergent functionality of the hippocampus, including effective navigation through time and space^8^ and the formation of episodic memories^9^. The hippocampus does not work in isolation and has extensive connections with other brain regions. Oscillations, and their synchrony or coherence, are believed to play an important role in coordinating the activity between the hippocampus and downstream regions^10–12^.

Despite extensive prior investigation of the neuronal populations in CA1, recent work suggests that some cell types still lack proper characterization^13^. Here, we report a novel cell population, which expresses neuronal nitric oxide synthase (nNOS), and has a number of unique properties. These cells are GABAergic, with both extensive local and long-range axons, and we therefore refer to them as LINCs: long-range inhibitory nNOS-expressing cells. Although LINCs express nNOS and possess long-range axons, they do not appear to simply be hippocampal versions of cortical NOS-type I cells, nor do they closely match any other previously characterized cell population, as detailed here. Despite being relatively few in number, LINCs can have a major impact on hippocampal function, oscillatory power and coherence.

## RESULTS

### Intersectional vector approach targets LINCs

The ability to identify, characterize, and manipulate LINCs rests on the use of a recently developed Cre- and Flp-dependent virus for the expression of eYFP-tagged ChR2 (AAV-DJ-hSyn-Con/Fon-hChR2(H134R)-eYFP-WPRE^14^), and mice expressing Cre in nNOS+ neurons and Flpe in Dlx5/6+ GABAergic cells (**Fig.1a,b**). This approach should limit expression of eYFP-tagged ChR2 to nNOS-expressing interneurons. Potentially due to the location of injection and the serotype used (DJ), this protocol surprisingly did not result in labeling of the expected hippocampal nNOS-expressing interneuron populations^15–18^. Instead, this approach resulted in labeling of a population of cells (i.e. LINCs) with unexpected morphologies (**Fig.1d,f,g**; **Extended Data Figs.1,2)**. LINCs could be labeled with either a dorsal, or a more ventral, CA1 injection, with labeled cells found along the anterior-posterior extent of the hippocampus at a considerable distance from the site of injection (**Fig.2a**; **Extended Data Fig.3**), suggestive of widespread processes. In contrast, viral injection into animals negative for Cre or Flpe did not result in opsin expression (**Fig.1e**). We further confirmed, through immunohistochemistry, that virally-labeled LINCs express nNOS, although this can be difficult to detect in some cells (**Fig.1c; Extended Data Fig.1**). Together these findings suggested that this vector-based approach labels a unique population of nNOS+ neurons, and warranted further investigation.

**Figure 1.**
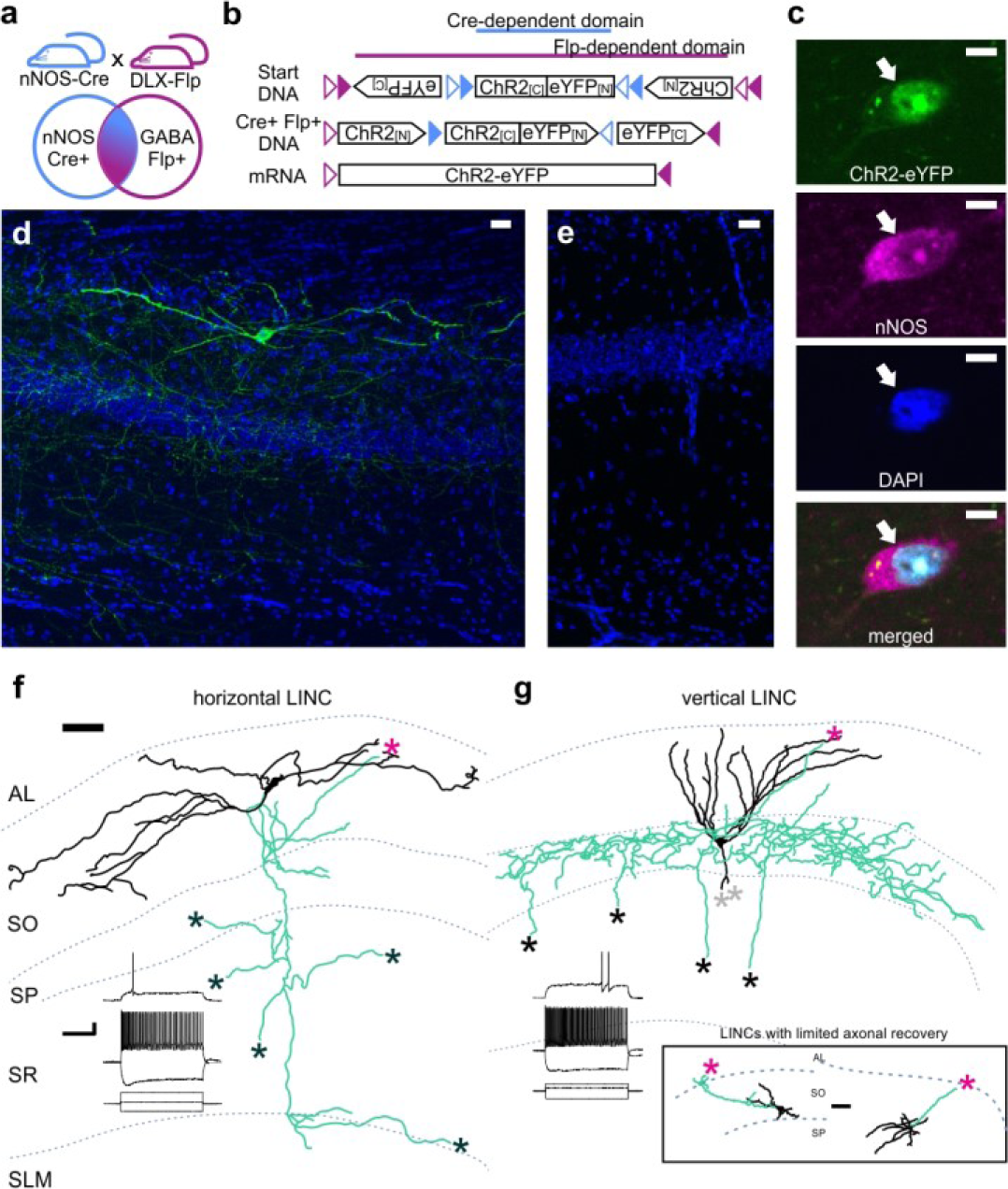
Intersectional genetic viral vector approach to target and characterize LINCs in the hippocampus. nNOS-Cre x Dlx5/6-Flpe mice (**a**) injected with an intersectional ChR2-eYFP AAV (**b**) show selective expression in nNOS-positive (**c**) inhibitory neurons (**d**, representative LINC). No expression is seen in injected negative control animals (**e**). LINCs have largely horizontal (**f**) or vertical (**g**) dendrites (black). In addition to local axons (green), all filled LINCs showed a severed (asterisks) axon, often en route to the alveus (magenta), present even when the axon was not well-recovered (inset box), suggestive of long-range projections. Insets: firing properties of the cells in **f, g.** Panel (**b**) based on ref [^14^]. Alveus (AL), stratum oriens (SO), stratum pyramidale (SP), stratum radiatum (SR), stratum lacunosum moleculare (SLM). Scale bars: 5μm (c); 25μm (**d-e**); 50μm (**f-g**); 20mV, 500pA and 200ms (**f-g**).

### LINCS provide broad & long-lasting inhibition

As our findings suggested that LINCs may have not been previously studied, we first sought to better characterize these cells by performing whole-cell patch clamp recordings from LINCs, to determine their electrophysiological properties and provide greater examination of their morphologies. Confirming observations based on eYFP-expression alone, recorded LINCs did not display electrophysiological nor morphological properties suggestive of previously characterized nNOS-expressing interneurons of CA1 **(Fig.1f,g; Extended Data Table 1**).

LINCs displayed either largely horizontal dendrites (hLINCs; **Fig.1f**), largely vertical dendrites (vLINCs; **Fig.1g**), or an intermediate ‘T-shaped’ morphology (**Fig.2d,e**). In both individually filled cells and in cells whose morphology was discerned from the expression of eYFP, we found that the dendritic morphology roughly corresponded to the somatic location: hLINCs were found primarily in the stratum oriens (SO), while vLINCs were found primarily in the stratum pyramidale (SP) (**Fig. 2e**). Previous literature does note the existence of nNOS-immunopositive cells in SO with “largely horizontal dendrites”, as well as occasional “bitufted” cells in SP, but any further characterization of these cells was lacking^2^.

**Figure 2.**
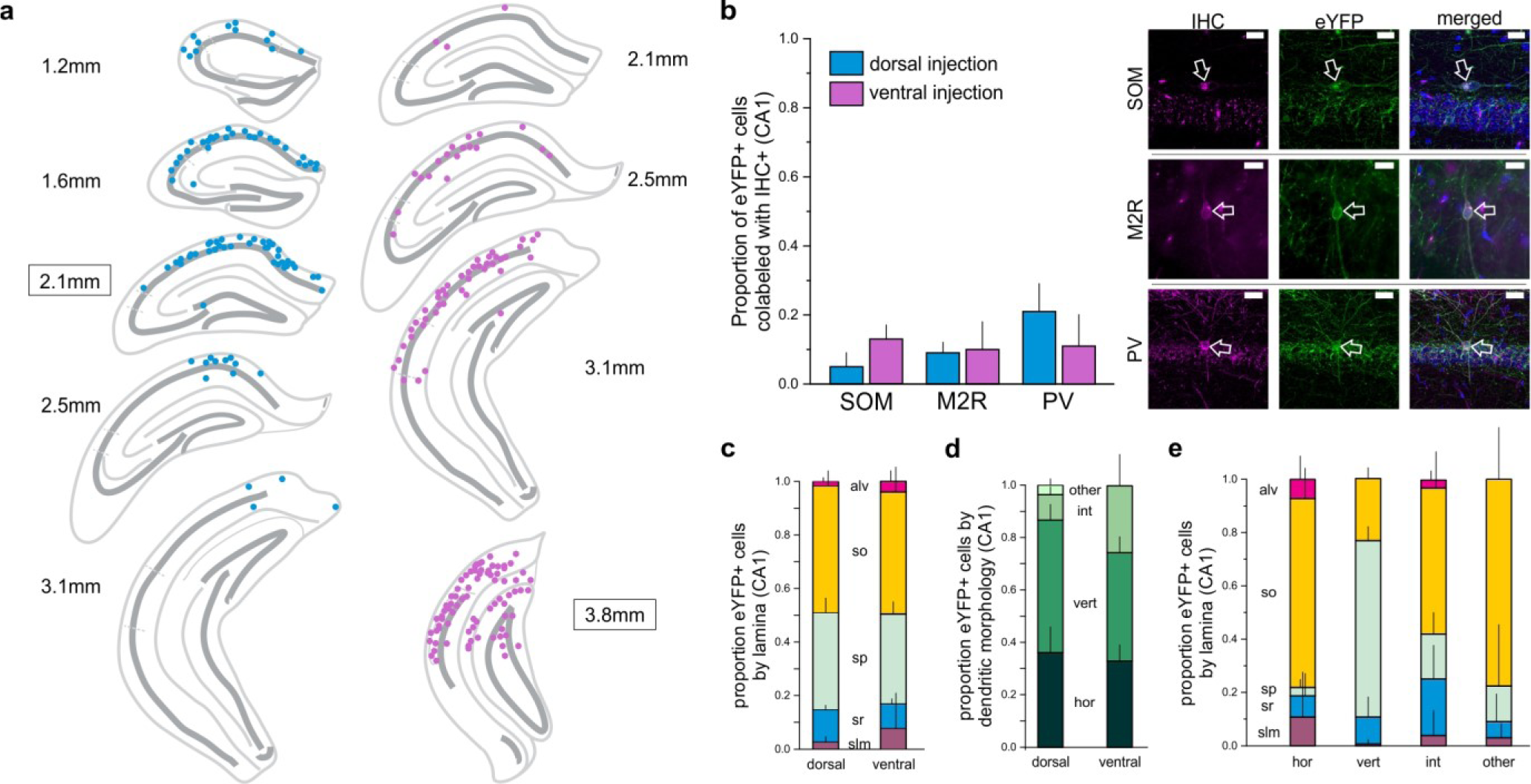
LINCs are located along the anterior-posterior extent of the hippocampus, and few express SOM, M2R, or PV. **a**) eYFP+ somata from a 1-in-4 coronal series after dorsal (blue) or more ventral (violet) virus injection (approximate injection position outlined). **b**) Somatostatin (SOM), muscarinic acetylcholine receptor 2 (M2R), or parvalbumin (PV) immunohistochemistry (IHC); n=3 animals (dorsal) and 3 animals (ventral) for each marker. DAPI in blue. **c-e**) CA1 LINCs by lamina (**c**), dendritic morphology (**d**), or both (**e**). Horizontal (hor), vertical (vert), intermediate (int), or other dendritic morphologies. Cells without discernable dendrites were counted as ‘other.’ (**b-e**) displays mean+SD. Scale bars=20μm (**b**).

Overall, LINCs had a modest input resistance (177 ± 17 MΩ), a threshold for firing near −44mV (−44.6 ± 0.5 mV), and a relatively low firing frequency near threshold (31 ± 6 Hz, mean±SEM, n=21 cells). LINCs also displayed subtle variability in their firing properties, which, to a limited extent, corresponded to dendritic morphology **(Fig.1f,g insets**; **Extended Data Table 1**). Specifically, maximum firing frequency (hLINCs: 149 ± 22 Hz; vLINCs: 78 ± 18 Hz; hLINC vs vLINC, uncorrected *p*=0.02, two-tailed Mann-Whitney (M-W) test, mean±SEM), adaptation ratio (hLINCs: 0.39 ± 0.05; vLINCs: 0.63 ± 0.05; hLINC vs vLINC uncorrected *p*=0.009, M-W) and coefficient of variance of the interspike interval (hLINCs: 11.7 ± 2.6; vLINCs: 24.4 ± 3.6; hLINC vs vLINC uncorrected *p*=0.009, M-W) were suggestive of differences, with hLINCs showing a slightly faster and more consistent rate of firing. **Extended Data Table 1** provides a detailed list of LINCs’ electrophysiological properties. Overall, LINCs had firing properties consistent with “regular spiking” or “regular spiking non-pyramidal” descriptions^19–21^.

Inhibitory neurons are typically categorized primarily by their axonal, rather than dendritic, arbor^2,3^. Regardless of slice orientation, many LINCs displayed poor axonal recovery, with the axon often lost as it ascended towards the alveus, suggestive of long-range projections^22,23^. Indeed, all filled LINCs displayed an axon that was lost as it exited the sectioned tissue (21/21 filled LINCs, **Fig.1f,g**, **Extended Data Fig.2**). For the cells in which a more extensive local axonal arbor was recovered, a variety of axonal arbors were found (**Fig.1f,g**), which did not appear to correspond strongly to dendritic morphology (**Extended Data Fig.2**). No drumstick-like appendages^23,24^ on the axons of LINCs were noted. Despite the generally poor axonal recovery from LINCs filled during *ex vivo* hippocampal recordings, we found that, collectively, local axons reached all layers of CA1 (**Fig.1**, **Extended Data Fig.2)**, suggesting a potentially broad impact of LINCs on neurons in the region. We therefore next asked what populations of CA1 cells are targeted by these local axons.

Previous work indicates heterogeneity in hippocampal pyramidal cells^25–28^, including their inhibition by local interneurons^28,29^. Therefore, to determine LINCs’ local connectivity, we recorded from both deep and superficial pyramidal cells, as well as inhibitory neurons across all layers of CA1, while optogenetically activating LINCs. We found that LINCs broadly targeted both deep and superficial CA1 pyramidal cells (dPC and sPC, respectively; **Fig.3**), to a roughly equivalent degree: GABA_A_ responses (subsequently blocked by 5μΜ gabazine) were recorded in approximately 80% of dPCs and sPCs (13/16 dPCs; 16/20 sPCs; *p*=0.93, χ^2^ test; **Fig.3a,d**), and were of similar amplitude in both dPCs and sPCs (median dPC: −106pA, median sPC: −83pA; dPCs vs sPCs, *p*=0.97, M-W; **Fig.3b**; including non-responders, median dPC: −87pA, sPC: −66pA, *p*=0.29, M-W). This indicates that LINCs provide broad inhibition to both deep and superficial pyramidal cells in CA1, and therefore would link these two information processing streams^30,31^. Notably, LINCs also displayed similarly broad targeting of inhibitory neurons (INs), with approximately 80% of recorded INs also showing a postsynaptic GABA_A_ response (26/32 INs; vs dPCs *p*>0.99, χ^2^; vs sPCs *p*=0.91, χ^2^; **Fig.3**; median: −114pA, INs vs dPC vs sPCs, *p*=0.88, Kruskal-Wallis ANOVA (K-W); median including non-responders: −86pA, IN vs dPC vs sPC, *p*=0.62, K-W), with slightly stronger inhibition provided to INs with somata in SP (**Extended Data Fig.4**). Taken together, these findings indicate that LINCs provide unusually broad inhibition to CA1 cells.

In addition to a postsynaptic GABA_A_ response, a remarkably large percentage of recorded cells also displayed a postsynaptic GABA_B_-mediated response (present in gabazine, blocked by subsequent application of 5μΜ CGP55845; note that no postsynaptic responses remained after application of these GABA receptor antagonists, **Fig.3a**, further confirming the specificity of ChR2 targeting to only GABAergic neurons). Approximately 30-50% of recorded cells displayed a postsynaptic GABA_B_-mediated response (8/16 dPCs, 6/20 sPCs, and 10/32 INs; dPCs vs sPCs *p*=0.22, dPCs vs INs *p*=0.24, sPCs vs INs *p*=0.87, χ^2^; **Fig.3d**), with roughly equal amplitude across cell groups (**Fig3c**; in cells with a response: median dPC: 17 ± 3 pA, median sPC: 28 ± 13 pA, and median IN: 38 ± 10 pA, dPC vs sPC vs INs *p*=0.14 K-W; including non-responders: 8 ± 3 pA in dPCs, 2 ± 6 pA in sPCs, 9 ± 6 pA in INs, *p*=0.28 K-W). Taken together, this data indicates that LINCS provide strong, broad inhibition (through GABA_A_-mediated inhibition) as well as long-lasting (GABA_B_-mediated) inhibition in CA1. This would place LINCs in an influential position, capable of having a major impact on hippocampal function.

**Figure 3.**
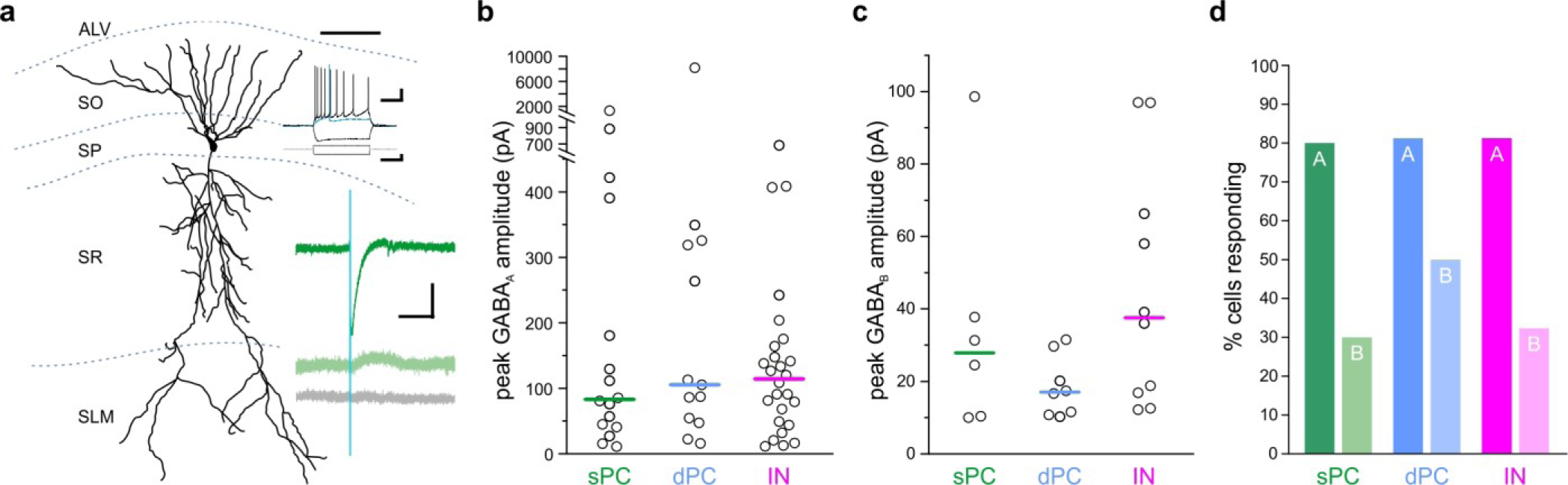
LINCs provide broad and long-lasting inhibition. Optogenetic activation of LINCs produces postsynaptic inhibitory responses in superficial pyramidal cells (sPC, green; example morphology, firing properties, and light-evoked inhibitory post-synaptic currents (IPSCs) shown in a), deep pyramidal cells (dPC, blue), and inhibitory neurons (IN, pink). **a**) Top green trace: light-evoked IPSC; middle green trace: in gabazine; bottom gray trace: in gabazine plus CGP55845. **b, c**) Peak amplitudes of GABA_A_ (**b**) or GABA**B** responses (**c**) in individual cells showing a response. Bar denotes median amplitude. **d**) Percentage of sPCs, dPCs, and INs with GABA_A_ response (denoted with ‘A’) or GABA_B_ response (‘B’). Scale bars (**a**): 50μm; 20mv, 200ms (top set); 400pA, 200ms (middle); 50pA, 200ms (bottom).

**Figure 4.**
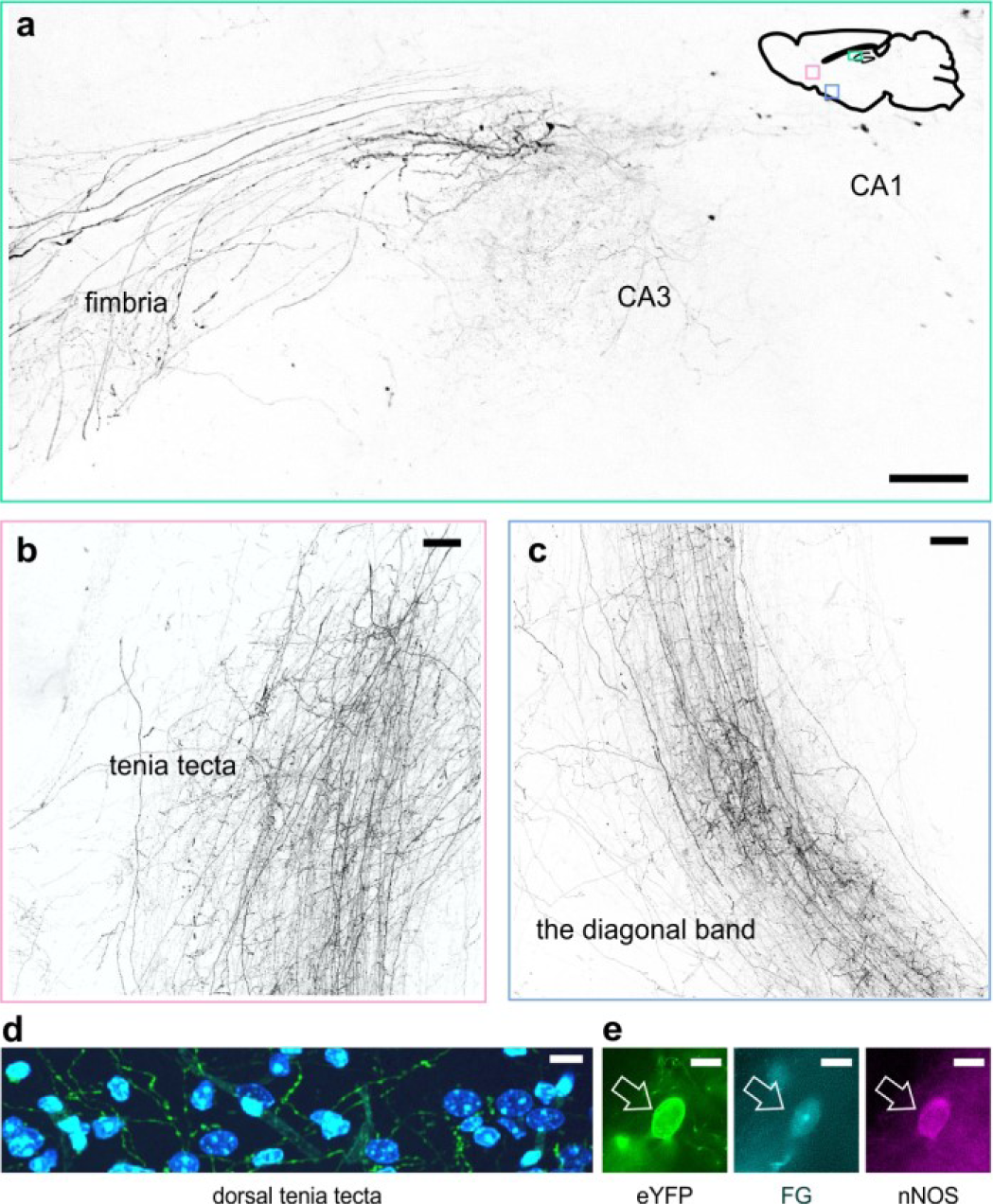
LINCs have extrahippocampal projections. LINCs project out of the hippocampus (**a**), through the medial septum (Extended Data Fig.4), and into the tenia tecta (**b, d**), the diagonal band (**c**), and other areas (Extended Data Fig.4). **a-c**) Max projections from X-CLARITY cleared tissue. d) eYFP+ processes in the dorsal tenia tecta, DAPI in blue. **e**) Example LINC colabeled with the retrograde tracer Fluorogold (FG) and confirmed nNOS IHC+ following injection of FG into the diagonal band. Scale bars: 100μm (**a-c**), 10μm (**d,e**).

**Figure 5.**
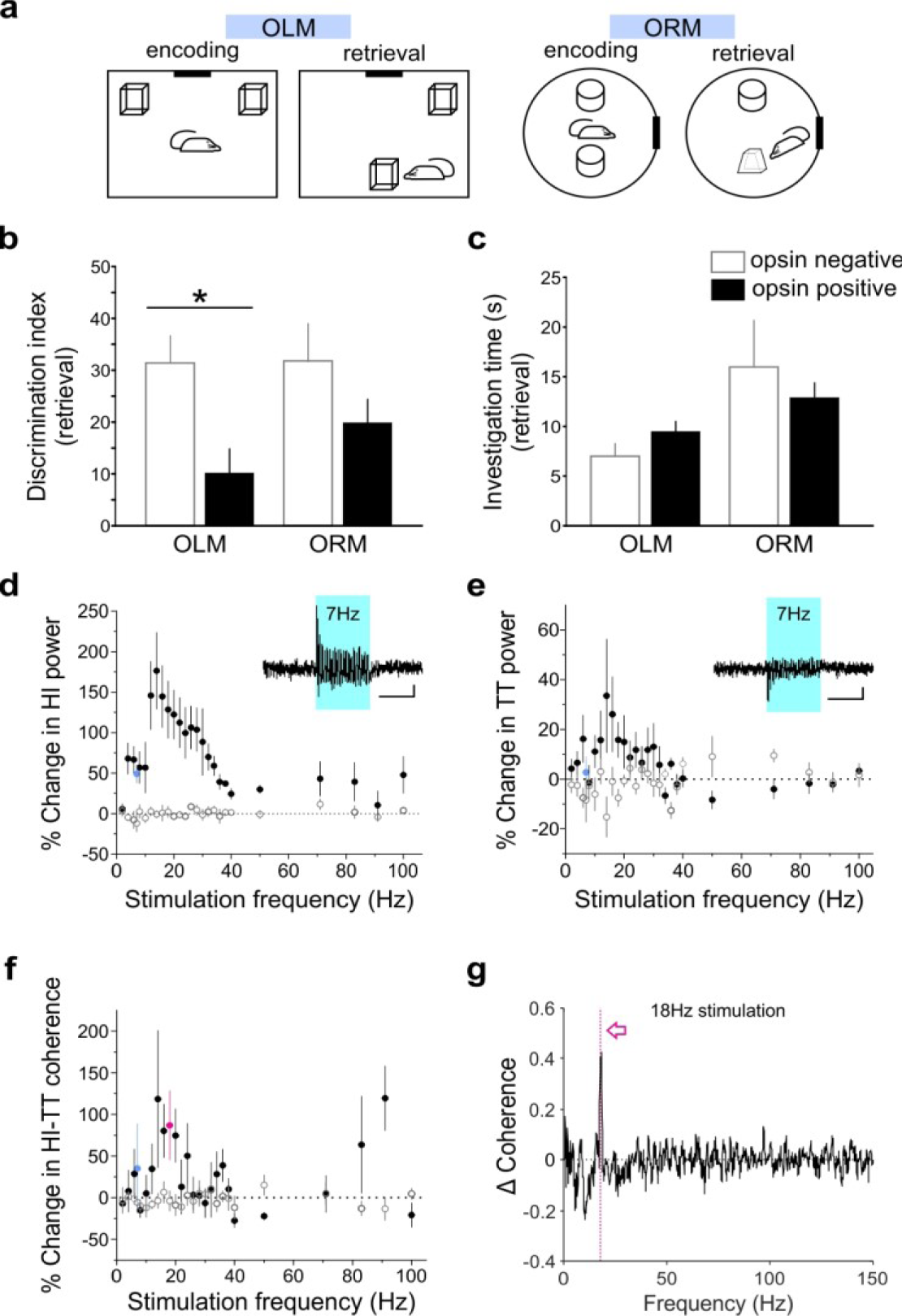
Manipulating LINCs *in vivo* affects spatial memory, oscillatory power, and hippocampal-frontal cortex coherence. **a**) Schematics for object location and recognition memory (OLM, ORM, respectively) tasks. **b**) Light stimulation significantly decreases performance (i.e. discrimination index) on the OLM task but not on the ORM task. **c**) Light stimulation does not affect total object investigation time during retrieval. **d-e**) Percent change in hippocampal (HI, **d**) and tenia tecta (TT, **e**) power at the stimulation frequency during light stimulation (opsin+: closed circles, opsin-negative: open circles). Insets: Light delivered to the hippocampus evokes responses in the hippocampus (**d**) and tenia tecta (**e**), mean traces from 120 light deliveries. **f**) Percent change in hippocampal-tenia tecta coherence at the stimulation frequency. **g**) Change in coherence in an opsin+ animal with 18Hz stimulation (magenta, **f**). **b-e**) Data shown: mean±SEM. Asterisk: *p-*value<0.05, M-W, opsin+ vs opsin-negative. Individual animals’ data shown in Extended Data Fig.6.

### LINCs have long-range projections

While the majority of hippocampal GABAergic neurons are true interneurons, with axons limited to targeting local neurons, there are notable exceptions, including cell-types that project far outside the hippocampus^32–34^. As noted above, recovered morphologies of LINCs recorded *ex vivo* were suggestive of long-range, extrahippocampal projections. We further examined this possibility in X-CLARITY cleared tissue, as well as traditionally sectioned tissue (**Fig.4**; **Extended Data Fig.5**). Fibers exiting the hippocampus through the fimbria were clearly visible (**Fig.4a**; **Extended Data Video 1**). These fibers continued down through the medial septum (**Extended Data Fig.5)**, and into the dorsal and ventral tenia tecta (**Fig.4b,d;** a frontal lobe brain region also receiving projections from CA1 PCs^35,36^) and the vertical and horizontal limbs of the diagonal band of Broca (**Fig.4c**; **Extended Data Fig.5)**. LINCs appear to provide broad connections from the hippocampus to other brain regions: labeled fibers were also identified in the dorsal subiculum, entorhinal cortex, mammillary nuclei, lateral hypothalamus, olfactory tubercle, olfactory bulb, ipsilateral dentate gyrus (with some fibers projecting through CA3 and others crossing the hippocampal fissure), and the contralateral hippocampal formation (including the contralateral dentate gyrus; fibers visible in the dorsal hippocampal commissure) (**Extended Data Fig.5**). To further confirm that CA1 LINCs project to extrahippocampal areas, we injected the retrograde tracer Fluorogold (FG) into the tenia tecta or diagonal band of Broca and observed co-labeling with eYFP and FG in CA1 LINCs (**Fig.4e**). Therefore, in addition to extensive targeting within CA1, LINCs provide long-range input to a variety of extrahippocampal brain regions, positioning them to also play a role in interregional communication or synchrony.

A previously identified population of GABAergic CA1 cells with extrahippocampal projections, referred to as hippocampal-septal (or double-projection^34^) cells, also reside in the SO of CA1^2,22,23,34,37–39^. To test the hypothesis that LINCs are actually hippocampal-septal or double-projection cells, we performed immunohistochemistry for somatostatin (SOM). While hippocampal-septal (and double-projection) cells express SOM at high rates (reported as 100% ^22,34^), only a small subset of LINCs were immunopositive for SOM (SOM+) in CA1 (4.9 ± 4.4%, n=3 animals, dorsal injection; 12.5 ± 4.4%, n=3 animals, ventral injection; **Fig.2b**, **Extended Data Fig.3**). SOM expression was also not highly enriched in the FG-labeled LINC population; no FG+ LINCs were SOM+ after FG injection into the tenia tecta (n=3 animals), and only 17.2 ± 14.1% of FG+ LINCs were SOM+ after FG injection into the diagonal band of Broca (4 of 29 FG+ LINCs in CA1, n=3 animals). Therefore, LINCs are a population of cells distinct from the previously described hippocampal-septal cells. Notably, this data further argues that LINCs are also not simply hippocampally-located versions of neocortical NOS-type I cells. Neocortical NOS-type I cells, like LINCs, have long-range projections and express nNOS^18,40^, but, unlike LINCs, NOS-type I cells also express SOM at high levels and rates^18,41^. Therefore, minimally, expression levels of SOM distinguish LINCs from NOS-type I cells. Similarly, LINCs are distinct from nNOS cells identified as likely corresponding to “backprojection” cells^42^ based on gene expression profiling, which have consistently high levels of SOM expression^13^. Consider also the lack of axonal drumstick-like appendages^23,24^ on LINCs (**Fig.1**, **Extended Data Fig.2**). Therefore, LINCs are distinct from these previously characterized long-range projecting GABAergic neurons.

High postsynaptic connectivity and long-range projections are reminiscent of early-generated (EG), GABAergic hub cells of CA3, which are capable of orchestrating network-wide synchronous activity^22,43,44^. Similar to LINCs, while hub cells are unified by their widespread axonal arborization, they display some morphological heterogeneity, including in both axonal structure (i.e., some hub cells are perisomatic targeting – compare to LINC in **Fig.1g** – while others have dendritically targeting axons – compare to LINC in **Fig.1f)**^43^ and dendritic morphology (including cells with largely horizontal or largely vertical dendrites)^44^. Additionally, both EG GABAergic hub cells and LINCs have broad hippocampal and extrahippocampal targets. However, LINCs also have notable differences from EG hub cells, including firing properties^44^ and expression levels of SOM (prevalent in EG hub cells) and nNOS (uncommon in EG hub cells)^44^.

Other hippocampal GABAergic projection neurons, including trilaminar cells and oriens-retrohippocampal cells are commonly associated with the muscarinic acetylcholine receptor 2 (M2R)^34,42^. We therefore also tested LINCs for this marker, but again found relatively few LINCs being immunolabeled (M2R-immunopositive CA1 LINCs: 8.5 ± 3.2%, n=3 dorsally injected animals; 9.7 ± 8.0%, n=3 ventrally injected animals; **Fig. 2b**; **Extended Data Fig.3**). Finally, we additionally tested for parvalbumin (PV), as previous work has shown that hippocampal PV+ cells can project to the contralateral hippocampus^45,46^. We found only limited labeling with this marker (PV-immunopositive CA1 LINCs: 20.9 ± 7.7% n=3 dorsal, 11.5 ± 9.3% n=3 ventral, **Fig.2b**; **Extended Data Fig.3**), consistent with the overall spike width and firing rates of LINCs (**Extended Data Table 1**). Taken together, it is evident that LINCs are best labeled with nNOS (although nNOS expression is not limited to LINCs^2,16,18^), rather than other canonical markers of hippocampal projecting inhibitory neurons, and that LINCs do not fit well into previously described CA1 GABAergic cell populations.

### LINCs impact hippocampal function

Despite being relatively sparse in number, LINCs have long-range projections suggesting a role in interregional communication, and provide broad local inhibition, suggesting an influential role in the hippocampus. We therefore next set out to test LINCs’ ability to influence hippocampal function and oscillations *in vivo.* First, we tested the hypothesis that manipulating LINC activity would strongly impact hippocampal function. To test this hypothesis, we optogenetically manipulated LINCs *in vivo* during the object location memory (OLM) task and the object recognition memory (ORM) task (**Fig.5a**). The OLM task is strongly hippocampal dependent^47,48^. In contrast, the ORM task, which nicely parallels the OLM task in format ^49^, is not strongly hippocampal dependent^47,50^.

Optogenetic manipulation of LINCs (3s of ~7Hz stimulation^51^ every 30s during encoding and retrieval) had no significant effect on the ORM (discrimination index opsin+: 19.9 ± 4.5, vs 31.7 ± 7.3 for opsin-negative, n=16 and n=8 animals, respectively, *p*=0.34, M-W; **Fig.5b**; **Extended Data Fig.6**), but produced strong spatial memory deficits in the OLM task (OLM discrimination index: 10.1 ± 4.7, n=14 opsin+ animals; vs 31.4 ± 5.2, n=8 opsin-negative animals, *p*=0.009, M-W; **Fig.5b**; **Extended Data Fig.6**). Confirming that these effects were not due to a general indifference to the objects nor to motor deficits, there was no effect on the total time exploring objects (OLM opsin+: 9.5 ± 1.0s, vs 7.0 ± 1.3s opsin-negative, *p*=0.16; ORM opsin+: 12.9 ± 1.5s, vs 16.0 ± 4.7s opsin-negative, *p*=0.98; **Fig.5c**; **Extended Data Fig.6)**. These findings illustrate that manipulation of hippocampal LINCs strongly impairs performance on a spatial memory task, and therefore indicate that, despite being relatively few in number, LINCs can have a substantial impact on hippocampal function.

### LINCs impact oscillations & HI-frontal cortex coherence

We next sought to determine the impact of optogenetic activation of LINCs on hippocampal and extrahippocampal network synchrony. In addition to testing the parameters used for LINC stimulation during behavioral testing (~7Hz; **Fig.5**), we tested a range of stimulation frequencies, to examine not only if oscillatory power can be altered by LINCs, but also which frequencies produce the greatest changes in hippocampal and extrahippocampal power or coherence. To do this, we simultaneously monitored the local field potential (LFP) from the hippocampus and the tenia tecta (TT; a frontal cortex brain region receiving input from CA1 LINCs; **Fig.4**), while optogenetically stimulating LINCs in the hippocampus. Stimulating LINCs in the hippocampus, at a variety of frequencies, strongly affected hippocampal power at that stimulation frequency (mixed-design ANOVA with Greenhouse-Geisser correction: genotype *p*=0.00008, F: 46, degrees of freedom (DF): 1; genotype*frequency *p*=0.00082; F: 6.6, DF: 3.5; **Fig.5d**; **Extended Data Fig.7**), with stimulation between roughly 4Hz and 40Hz showing the greatest entrainment of the LFP to the stimulation frequency. A smaller change in power at the stimulation frequency was apparent in the TT (two-way repeated measures ANOVA with Greenhouse-Geisser correction: location *p*=0.0012, F: 66.5, DF: 1; location*frequency *p* = 0.018, F: 4.7, DF:3.25; **Fig.5e** and **Extended Data Fig.7**).

Beyond increases in oscillation power, increases in oscillation synchrony (i.e. coherence) are believed to play an important role in coordinating activity between brain regions^10,11,52^. LINCs, having connectivity both within the hippocampus and to extrahippocampal regions, are in a prime position to increase interregional coherence (i.e., to ‘link’ them up). Indeed, we measured significant increases in coherence between the hippocampus and TT when optogenetically activating LINCs, across a range of stimulation frequencies (mixed-design ANOVA with Greenhouse-Geisser correction: genotype *p*=0.0017, F:19.2, DF: 1; genotype*frequency *p*=0.14, F: 1.9, DF: 3.5; **Fig.5f,g**). Together, these data suggest that LINCs can impact hippocampal function, strongly entrain hippocampal oscillations, and increase coherence between the hippocampus and downstream regions.

## DISCUSSION

Here, we have taken advantage of recent advances in viral vector specificity (i.e. the INTRSECT^14^ approach) to selectively label and manipulate a population of CA1 cells previously lacking any detailed description: LINCs. In addition to this being the first time LINCs have been described in any detail, this work revealed that LINCs have several properties that make them especially unique and exciting: 1) LINCs have widespread postsynaptic connections within the hippocampus, allowing them to provide both fast and long-lasting GABAergic inhibition to almost any cell in CA1. 2) LINCs have long-range projections to many distinct regions of the brain. 3) Manipulating LINCs can cause spatial memory deficits, indicating that LINCs can have a significant impact on hippocampal function, and 4) LINCs can drive oscillatory activity in the hippocampus and increase interregional coherence. In summary, LINCs are poised to have a significant impact on the network, and a detailed understanding of LINCs is therefore important for proper understanding of the hippocampal formation, and its downstream connections, more broadly.

Given the extensive prior examination of inhibitory neurons in CA1^2,3^, it seems surprising that any cell population, especially one with such widespread connections as LINCs, would have evaded characterization. In this regard, it is important to consider that nNOS expressing cells in the SO and SP with dendrites suggestive of LINCs had been noted^2^, but that further investigation was hampered. Indeed, many different factors have likely contributed to the prior difficulty in studying these cells. First, nNOS immunohistochemistry is notoriously challenging^53^, and LINCs can express relatively low levels, as well as dendritically-concentrated nNOS^53^, further complicating easy detection **(Extended Data Fig.1)**. Moreover, we found that other common long-range projection molecular markers are insufficient for labeling LINCs (**Fig.2**). Additionally, as nNOS is expressed in other CA1 populations, including pyramidal cells^53^, a simple nNOS-Cre based approach to targeting these cells would be insufficient (i.e., the recently developed intersectional approach^14^ is key). Indeed, our selective labeling of LINCs was also due to serendipity (and likely the AAV serotype used), as other interneuron populations also express nNOS^2,16,18^. Similarly, retrograde-based labeling or expression systems suffer from the relative rarity of LINCs and the fact that many of the areas targeted by LINCs are also targeted by other cell populations – for example, CA1 pyramidal cells target the TT^35,36^ which may have overwhelmed the ability to previously identify LINCs, especially those residing in the SP^23^. In light of these numerous complications, it is not entirely surprisingly that LINCs have largely evaded previous consideration.

Our data clearly indicate that LINCs should no longer be overlooked; they have a broad impact on the hippocampus, and appear to play a role in coordinating hippocampal and extrahippocampal activity. Future work will provide important additional information about the roles of LINCs not only in healthy physiology, but also pathophysiology, and if manipulation of LINCs could provide therapeutic benefit, for example, in Alzheimer’s disease^54^ or temporal lobe epilepsy^51^.

## METHODS

All experimental protocols were approved by the University of Minnesota’s Institutional Animal Care and Use Committee.

### Animals

For all experiments, mice were bred in-house. Dlx5/6-Flpe founders, expressing Flpe recombinase in GABAergic forebrain neurons, and RCE:dual founders were kindly provided by the Fishell lab (also available from Jackson Laboratory (fDLX: Tg(ml56i-flpe)39Fsh/J, stock 010815, maintained as hemizygotes^55^; RCE:dual: Gt(ROSA)26Sor^tm1CAG-EGFP)Fsh^/Mmjax, stock 32036-JAX, maintained as hemizygotes^55^). nNOS-Cre founders, expressing Cre recombinase in nNOS-expressing neurons, were purchased from Jackson Laboratory (cNOS; B6.129-Nos1^tm1(cre)Mgmj^/J; stock 017526; maintained as homozygotes^56^). fDLX mice were crossed with cNOS mice to produce cNOS-fDLX mice, which were used for experiments. Positive offspring expressed both Cre and Flp recombinases in nNOS-expressing GABAergic neurons of the forebrain; Flpe-negative littermates were used as controls. Uncrossed Flpe+ fDLX mice (n=2) and Black6 (n=2, C57BL/6J, Jackson Laboratory stock 000664) mice were injected with virus to further confirm specificity of viral expression in the absence of Cre. Both male and female mice were used in experiments; no significant differences were noted between males and females regarding properties of LINCs, post-synaptic responses, discrimination indices, nor changes in power or coherence, and sexes have been combined throughout. Except following implantation, animals were housed in standard housing conditions in the animal facility at the University of Minnesota. Following implantation for behavioral experiments and *in vivo* electrophysiology, animals were singly housed in investigator managed housing. In all housing conditions, animals were allowed *ad libitum* access to food and water.

### Stereotaxic Surgeries

Surgical procedures were performed stereotaxically under isoflurane anesthesia^45,51,57,58^.

#### Viral & Fluorogold Injections

cNOS-fDLX mice were injected with 1μL of a virus encoding the excitatory opsin channelrhodopsin (hChR2(H134R); ChR2) fused to enhanced yellow fluorescent protein (eYFP) in a Cre- and Flp-dependent manner (AAV-DJ-hSyn-Con/Fon-hChR2(H134R)-eYFP-WPRE ^14^; UNC Viral Vector Core lot numbers AV6214 and AV6214C, titer 4.4×10^12^ vg/mL; Stanford Viral Vector Core lot numbers 1599, 3214, titers 2.0×10^13^ and 2.55×10^13^ vg/mL respectively) via Hamilton syringe (model 7002KH) into the left dorsal hippocampus (0.2cm posterior, 0.125cm left, and 0.16cm ventral to bregma) or ventral hippocampus (0.36cm posterior, 0.28cm left, and 0.28cm ventral to bregma) at an approximate rate of 200nL/min at postnatal day 45 or greater. After the full volume of virus was injected, the syringe was left in place for at least 5 minutes before being withdrawn. Animals recovered on a heating pad and were returned to the animal housing facility the following day. Experiments were conducted at least 6 weeks following viral injection. Fluorogold (FG; 100nL, 4% in saline; Fluorochrome LLC, cat#52-9400) was similarly injected into the tenia tecta (0.22cm anterior, 0.05cm left, and 0.375cm ventral to bregma) or vertical limb of the diagonal band (0.1cm anterior and 0.5cm ventral to bregma). For each day of FG injections, one FG-injected brain was harvested acutely to confirm targeting of the tracer injection (n=2 acute brains). The remaining FG-injected brains (n=3 tenia tecta, 3 diagonal band) were harvested 1 week following FG injection.

#### Implant Surgery

Mice used for behavioral and *in vivo* electrophysiology experiments were additionally implanted with a twisted-wire bipolar electrode (PlasticsOne, 2-channel stainless steel, MS303/3-A/SPC) and optical fiber (Thorlabs, FT200UMT, Ø200μm, 0.39 NA) in the dorsal hippocampus near the injection site (0.2cm posterior, 0.125cm left, and 0.15cm ventral to bregma). A second twisted-wire bipolar electrode was implanted in the tenia tecta. Experiments were conducted minimally 5 days following the implant surgery to allow for recovery.

### Slice Electrophysiology Recordings

cNOS-fDLX mice previously injected with virus were deeply anesthetized with 5% isoflurane and their brains were dissected. Coronal or sagittal hippocampal sections were prepared in ice-cold sucrose solution and incubated at 36°C for 1 hour before being adjusted to room temperature until recording. All recordings were done at physiological temperature in artificial cerebrospinal fluid (ACSF). The sucrose solution contained the following (in mM): 85 NaCl, 75 sucrose, 2.5 KCl, 25 glucose, 1.25 NaH_2_PO_4_, 4 MgCl_2_, 0.5 CaCl_2_, and 24 NaHCO_3_. The ACSF solution contained (in mM): 2.5 KCl, 10 glucose, 126 NaCl, 1.25 NaH_2_PO_4_, 2 MgCl_2_, 2 CaCl, and 26 NaHCO_3_.

Slices were visualized with an upright microscope (Nikon Eclipse FN1) equipped with a xenon light source for visualizing fluorescence and optogenetic experiments (Lambda DG-4Plus, Sutter Instrument Company, model PE300BFA). Recordings were performed using pipettes (3-4ΜΩ) filled with an intracellular solution with a relatively high chloride concentration to record GABA_A_-mediated currents, and cesium-free, to allow recording of GABA_B_–mediated currents; the intracellular solution contained the following (in mM): 90 potassium gluconate, 43.5 KCl, 1.8 NaCl, 1.7 MgCl_2_, 0.05 EGTA, 10 HEPES, 2 Mg-ATP, 0.4 Na2-GTP, 10 phosphocreatine, and 8 biocytin, pH 7.29, 290 mOsm.

LINCs were quickly identified for recordings based on their eYFP fluorescence and were later confirmed to be opsin-expressing based on their light response that was resistant to antagonists. When determining LINCs’ postsynaptic targets, cells were pseudo-randomly patched and were *post hoc* confirmed eYFP-negative. For all recordings, after establishing whole-cell configuration, the resting membrane potential was immediately recorded and the firing pattern of the recorded cells was examined from a resting membrane potential adjusted to be approximately −60mV.

Recorded cells were then voltage clamped at −60mV and tested for a response to blue light (5ms duration, 10s intersweep interval) and the series resistance was monitored. The amplitude of the averaged postsynaptic response was measured at its peak (for GABA_A_ responses average time to peak: 6.3ms after the light was applied; for GABA_B_ responses: 134.0ms after light). A successful postsynaptic GABA_A_ response was defined as an inward current greater than or equal to 10pA below baseline; a GABA_B_ response was defined as an outward current greater than or equal to 10pA above baseline. Once a postsynaptic response was recorded, the GABA_A_ receptor antagonist gabazine (5μM, Sigma cat#S106) was bath applied and if a GABA_B_ receptor response remained, the GABA_B_ receptor antagonist CGP 55845 (5μM, Sigma cat#SML0594) was added to the bath. No response was ever evident after the application of CGP 55845.

After recordings, slices were fixed in 0.1M phosphate buffer with 4% paraformaldehyde for roughly 24 hours at 4°C. Biocytin filling was then revealed with Alexa Fluor 594-conjugated streptavidin (Jackson Immuno Research, 016-580-084, 1:500). Some sections were further processed for diaminobenzidine (DAB) and/or *camera lucida* morphological reconstructions. Cell identity (i.e. LINC, sPC, dPC, IN) was determined *post hoc* based on firing patterns, morphology, cell body location, and presence or absence of eYFP fluorescence.

### Immunohistochemistry, Tissue Processing & Cell Counting

Mice were heavily anesthetized with 5% isoflurane and decapitated. The brains were dissected and drop-fixed in 4% paraformaldehyde for roughly 48 hours, or, for some nNOS immunohistochemistry (**Extended Data Fig.1**), brains were drop-fixed in 1% paraformaldehyde for less than 24 hours^53^. Using a vibratome (Leica VT1000S), 50μm coronal or sagittal brain sections were collected in 0.1M phosphate buffer at room temperature. After fixation and sectioning, free-floating immunostaining was performed on every fourth section for either nNOS (rabbit anti-nNOS, Cayman Chemical, cat#160870, 1:1000), PV (rabbit anti-PV, Swant, PV27, 1:1000), SOM (rat anti-SOM, Millipore Sigma, MAB354, 1:250), or M2R (rabbit anti-M2R, EMD Millipore, AB5166, 1:1000), in red (goat anti-rabbit Alexa Fluor 594, Jackson Immuno Research, 111-585-003, 1:500; donkey anti-rat Alexa Fluor 594, Jackson Immuno Research, 712-585-153, 1:500). Sections were then mounted with Vectashield mounting media. Mounting media with DAPI was used for all tissue except those used in Fluorogold experiments.

Sections were visualized with epifluorescence or conventional transmitted light microscopy (Leica DM2500). eYFP-positive cells in the hippocampus were counted manually in every fourth 50μm section in all planes of focus. Once an eYFP-positive cell body was identified, its dendritic morphology was noted (horizontal, vertical, intermediate, or other) on the brain atlas and its soma was checked for colocalization with immunofluorescence and/or the retrograde tracer FG. In some instances, nNOS immuno-colocalization was found outside the soma in cells with nNOS immuno-negative cell bodies (**Extended Data Fig. 1**). Confocal imaging was performed on an Olympus FluoView FV1000 BX2 upright confocal microscope. Images in figures were adjusted for brightness and contrast, with all adjustments applied to the entire image.

### Tissue Clearing & Imaging

Animals (n=2) were perfused with ice cold phosphate buffered saline (PBS) followed by ice cold 4% PFA fixative, trimmed of excess right hemisphere tissue, and incubated in 4% PFA at 4°C overnight before active clearing^59^ with X-CLARITY (Logos Biosystems). Cleared samples were incubated in refractive index matched solution (RIMS)^60^ modified to preserve eYFP fluorescence and were imaged on a Nikon A1R inverted confocal microscope with a 10x glycerol immersion objective (0.5 numerical aperture, 5.5mm working distance) at the University of Minnesota’s University Imaging Centers.

### Behavioral Experiments

Object Recognition Memory (ORM) and Objection Location Memory (OLM) testing was performed as previously described [^49,57,61^] with minor modifications. In addition to the ORM and OLM tests, subjects underwent an additional task which was similar to the ORM, but odorants are presented on cotton swabs as a way to test odor recognition memory (OdorRM, modified after ref. [^62^]). Odorants (1-Octanol (0.1μL/mL), Sigma-Aldrich 95446, floral/citrus smell), nutmeg (1mg/mL, Target Market Pantry), and vanillin (0.2mg/mL, Sigma-Aldrich V1104) were diluted in mineral oil and were counterbalanced across mice (time spent investigating during encoding: 1-octanol: 5.7 ± 2.3 s, nutmeg: 5.4 ± 2.2 s, vanillin: 5.6 ± 2.2 s, mean ± SD). Each test took place over three days, with four days in between each round of testing. The order of the tests was counterbalanced across cohorts. For all tests, mice were habituated to the testing arena for 10 minutes on the first day. For the OdorRM test, two cotton swabs dipped in mineral oil were present throughout habituation. On the following (training/encoding) day, two identical objects (or two cotton swabs scented with the same odorant) were introduced to the arena and mice were allowed to explore for 10 minutes. On the third (testing/retrieval) day, two objects (or cotton swabs) were again placed in the arenas and mice were allowed to explore for 5 minutes. For the OLM, the objects remained the same on the testing/retrieval day, and one of the objects was moved to a new location (**Fig.5a**). For the ORM/OdorRM tests, the location of objects/odors was constant, but one of the objects/odors was replaced with a new object/odor (**Fig.5a**).

A train of blue light (5.0 ± 2.8 mW, Plexon PlexBright blue LED #94002-002 and driver #51382) pulses at approximately 7Hz frequency (50ms on, 100ms off; as in ref. [^51^]) was delivered for 3 seconds through the optical patch cable to the dorsal hippocampus once every 30 seconds during training and testing periods. Training and testing sessions were video-recorded and manually analyzed for time spent exploring each of the two objects/cotton swabs by an experimenter blinded to mouse genotype. Object/odor investigation time was defined in the videos as time when the mouse’s nose was within 1cm of the object/cotton swab’s edge and oriented towards the object/cotton swab, excluding time when the mouse was engaged in non-investigative behaviors (e.g. grooming or digging). A discrimination index (DI) was calculated for the testing day by subtracting the time spent investigating the familiar object/odor from the time spent investigating the novel object/odor and dividing the result by the total time investigating both objects/odors. Animals with a DI of greater than +/-20 on the encoding day were excluded (n=6 animals) as well as animals that investigated the objects for less than 3s (ORM/OLM, n=1 animal) or 2s (OdorRM, n=5 animals) during either training or testing. Note that animals displayed relatively low investigation times during the OdorRM task (**Extended Data Fig.6**), and the exclusion criteria for that task was adjusted to 2s accordingly. Additionally, n=2 positive animals were excluded for a lack of expression of the virus and n=2 animals were excluded for viral expression in the dentate gyrus rather than CA1.

### In vivo Electrophysiology Recordings

After behavioral testing, electrical patch cables were connected to the hippocampal and tenia tecta electrode pedestals. The electrical LFP signal (the local differential between the tips of the twisted wires of the electrode) was amplified 7,500-10,000 times (Brownlee Precision 410, Neurophase). A series of blue light stimulations was delivered to the dorsal hippocampus including the stimulation parameters used during behavioral training/testing as well as frequencies ranging from 2Hz to 100Hz (delivered in randomly assigned order, 30s intertrial interval, light pulse width 5ms, except for 7Hz stimulation which matched the light stimulation protocol from behavioral testing).

### Statistical Analysis

Statistical analyses were performed in MatLab R2014b and 2018a (including the Matlab Statistical Toolbox), and OriginPro 2016. A *p* value of less than 0.05 was considered significant. Data is shown as mean ± standard deviation (SD) unless otherwise specified.

#### Ex vivo electrophysiological recordings

LINCs’ electrophysiological properties were compared across section orientation (n=5 coronal, 18 sagittal), animal sex (n=15 female, 8 male), virus injection location (n=13 dorsal, 10 ventral/posterior), morphology of dendrites (n=11 horizontal, 10 vertical; 2 cells’ dendrites were not recovered and were excluded from this analysis), and cell body location (n=16 in the stratum oriens/alveus, 7 in the stratum pyramidale) using Mann-Whitney tests. There were no significant differences between section orientation, sex, or virus injection location, and these variables were collapsed in further analyses. Potential differences between LINCs with different dendritic morphologies were noted (**Extended Data Table 1** provides Bonferroni corrected alpha).

Postsynaptic response amplitudes were compared across cell types using a Kruskal-Wallis ANOVA (K-W) and *post hoc* two-tailed Mann-Whitney (M-W) tests when appropriate. Proportions of cell types showing postsynaptic GABA_A_ and/or GABA_B_ receptor-mediated responses and proportion of LINCs displaying persistent firing properties were compared using χ^2^ tests.

#### Behavioral testing

Retrieval discrimination indices (investigation_novel_ −investigation_familiar_)/(investigation_novel_ + investigation_familiar_) and total time spent investigating during retrieval were compared across genotype for each behavioral task using two-tailed Mann-Whitney tests.

#### In vivo oscillations & coherence

*In vivo* electrophysiological recordings were analyzed in MatLab 2018a with custom software utilizing the Chronux library (Chronux 2.12)^63,64^. For each animal and each light-delivery frequency, traces (sampled at 1000Hz) for both the hippocampal and the tenia tecta locations were aligned to onset of light delivery. The first step in analysis was to then remove traces which were likely to contain movement artifact: traces with a range (defined as the maximum recorded voltage value in the trace minus the minimum value) greater than two times the average range of all traces for that animal at that light-delivery frequency and location were removed prior to further analysis, leaving on average approximately 110 traces (median: 118 traces opsin+, 107 traces opsin-negative) for each animal (n=5 opsin+, n=6 opsin-negative animals) at each light-delivery frequency (26 stimulation frequencies).

To calculate the percent change in power at the stimulation frequency for each stimulation frequency for each animal, first the bandpower at the stimulation frequency (using a one hertz band centered around the stimulation frequency) was calculated for both the 3s prior to light-delivery (i.e. baseline) and the 3s during light delivery for each trial. These values were then averaged across trials, and the power during light expressed as a percent increase over baseline for each mouse at each stimulation frequency by recording location (**Fig.5d,e**; **Extended Data Fig.7**).

The Chronux library was similarly used to calculate the trial averaged coherence between the two locations (hippocampus and tenia tecta), both for the 3s prior to light delivery (baseline) and 3s during light delivery. Increase in mean coherence at the stimulation frequency (±0.5Hz) was then expressed as a percent increase from baseline (**Fig.5f**).

To visualize the percent change in power across frequencies during light delivery per location, genotype, and stimulation frequency, differential (i.e., percent increase) spectrograms **(Extended Data Fig.7)** were created as follows. For this analysis, frequencies plotted/analyzed were limited to up to 55Hz. Using Chronux, a trial averaged moving time spectogram was created for each animal/light-delivery combination for the 3s prior to light delivery (baseline), using a moving window size of 1s and a step size of 0.1s. The calculated power per frequency bin was then averaged across the 3s baseline time period. These averaged baseline values were then used to calculate the percent increase for the 3s of light stimulation (again, using a 1s window and 0.1s step size). The resulting spectrograms were then averaged across animals according to genotype, by recording location (displayed in **Extended Data Fig.7**).

### Data Availability

Data or custom code will be made available upon reasonable request.

---

**Extended Data Table 1.** Electrophysiological properties of LINCs by dendritic morphology. Two-tailed Mann-Whitney tests performed with Bonferroni corrected α; χ^2^ test performed for comparison of proportions showing persistent firing. Note that persistent firing^17,65^, also known as axonal barrage firing^66^, is associated with a different population of nNOS-expressing CA1 interneuron (i.e., 80% of neurogliaform cells display persistent firing, whereas only ~20% of other interneuron types display this phenomenon^17,66,67^), but is only rarely found in LINCs. Note also that despite potential subtle differences in firing pattern, LINCs show similar thresholds, input resistance, and firing frequency near threshold regardless of dendritic morphology.

| | Horizontal<br>(mean ± SEM) | Vertical<br>(mean ± SEM) | *p*-values<br>(corrected $\alpha$= 0.0045) |
| --- | --- | --- | --- |
| **Vrest (mV)** | -55.1 ± 2.5 | -59.7 ± 3.1 | 0.35 |
| **Input resistance (MΩ)** | 169 ± 20 | 191 ± 30 | 0.62 |
| **Threshold voltage (mV)** | -44.7 ± 0.6 | -44.2 ± 1.0 | 0.52 |
| **Firing frequency near threshold (Hz)** | 33 ± 11 | 30 ± 8 | 0.85 |
| **Max firing frequency (Hz)** | 149 ± 22 | 78 ± 18 | 0.02 |
| **Adaptation ratio of the ISI at max firing** | 0.39 ± 0.05 | 0.63 ± 0.05 | 0.009 |
| **Spike amplitude at threshold (mV)** | 54.4 ± 2.8 | 52.9 ± 1.5 | 0.62 |
| **Spike amplitude at max (mV)** | 39.7 ± 2.9 | 41.3 ± 3.0 | 0.43 |
| **Action potential half width at threshold (ms)** | 0.67 ± 0.10 | 0.96 ± 0.15 | 0.16 |
| **Action potential half width at max firing frequency (ms)** | 0.72 ± 0.15 | 1.20 ± 0.19 | 0.57 |
| **Coefficient of variance of the ISI at max firing** | 11.7 ± 2.6 | 24.4 ± 3.6 | 0.009 |
| **Proportion of cells showing persistent firing** | 0/9 | 2/10 | 0.16 |

**Extended Video 1.**
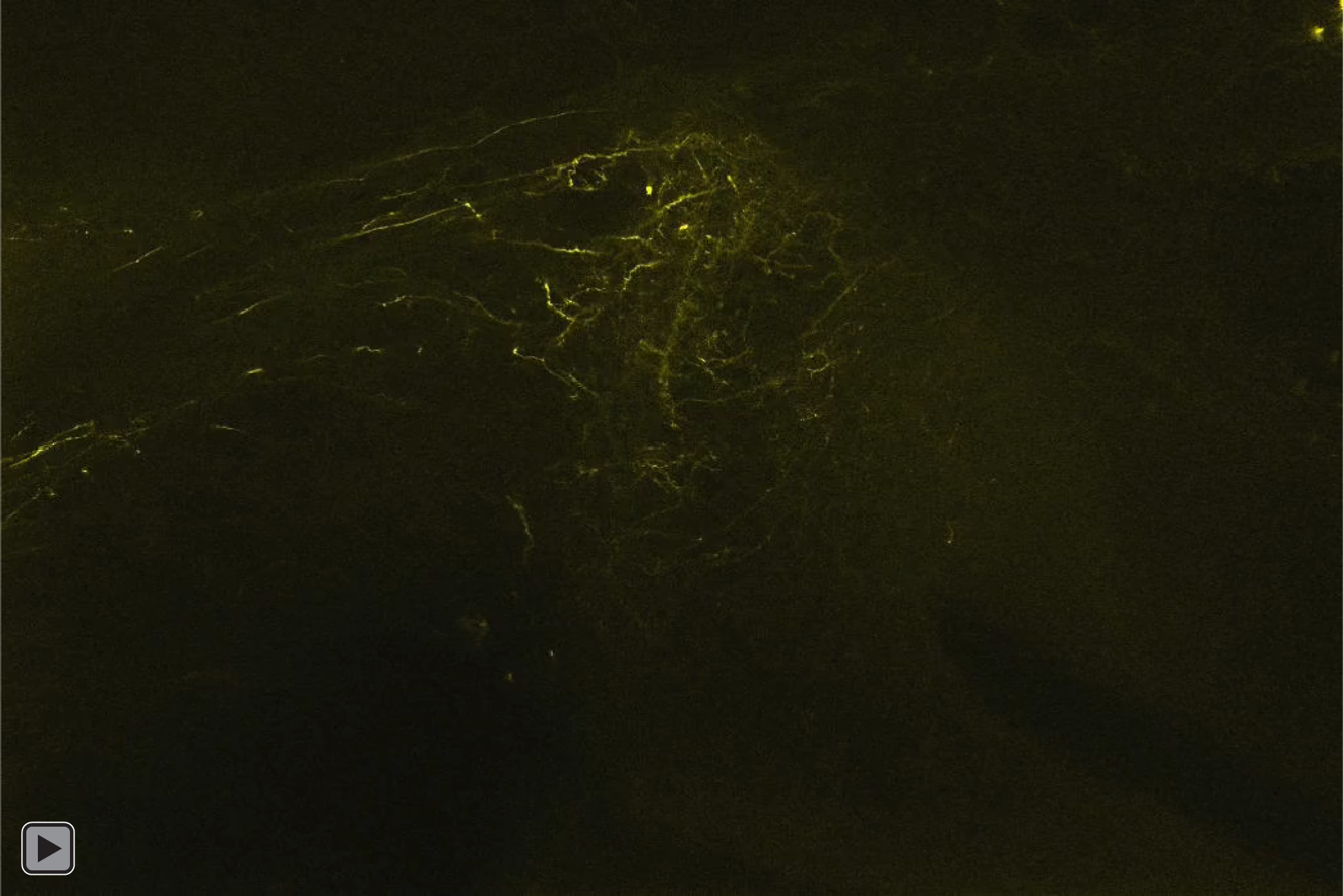
eYFP+ processes exiting hippocampus through fimbria in cleared whole brain tissue.

**Extended Data Figure 1.**
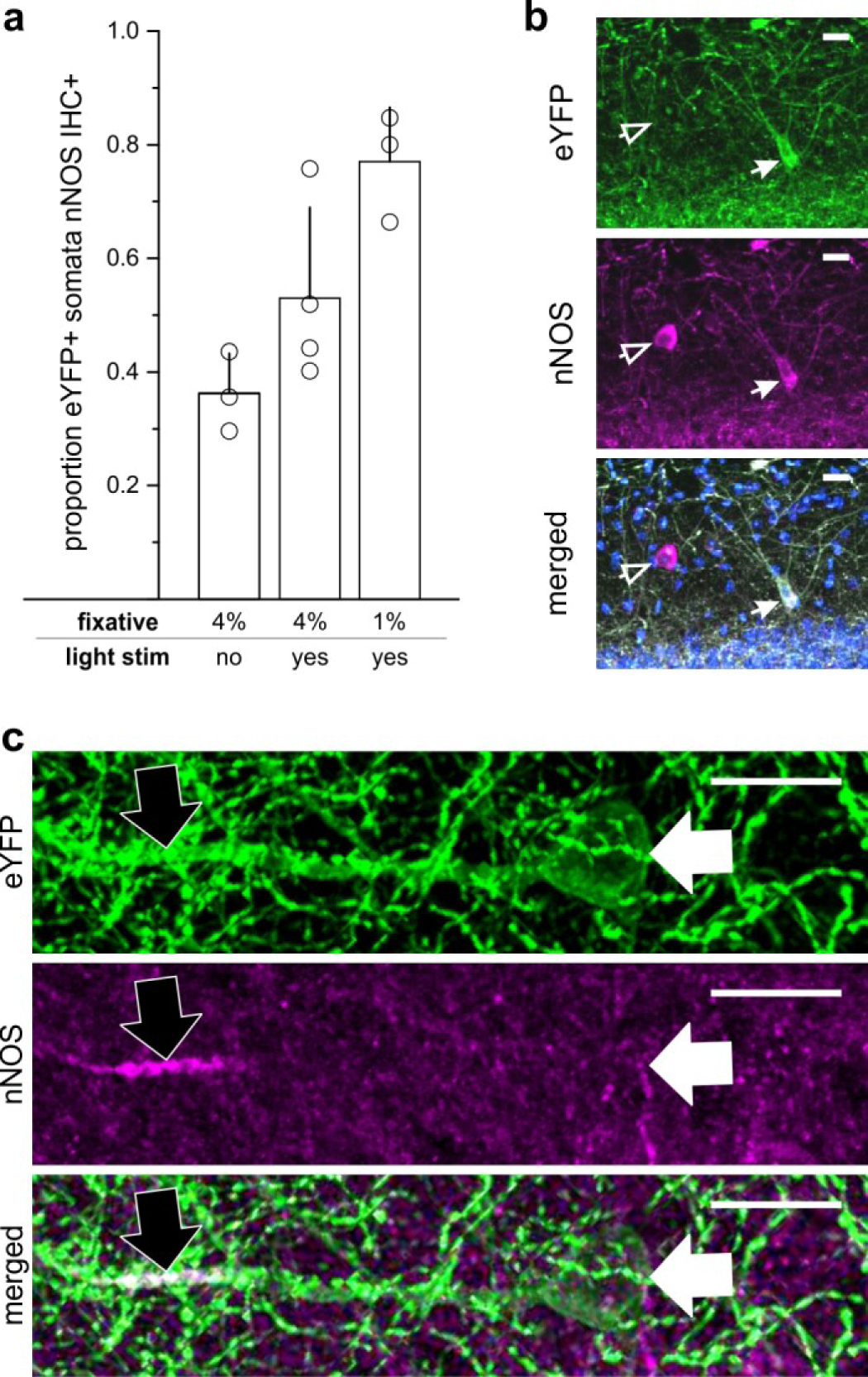
LINCs express nNOS. **a)** The proportion of eYFP+ somata that were nNOS IHC+ by experimental condition and percent paraformaldehyde (PFA) fixative. Without optogenetic activation of LINCs, and following 4% paraformaldehyde fixation, a limited number of LINCs displayed obvious somatic nNOS immunoreactivity (303/811 eYFP+ cells, 36.2 ± 7.0%, n=3 animals). While regulated expression of NOS is most strongly associated with iNOS, regulated expression of nNOS is also reported^68–71^. We therefore additionally examined the proportion of eYFP+ somata colocalizing with nNOS IHC after light delivery to optogenetically activate LINCs (4% PFA; 326/565 eYFP+ cells, 53.0 ± 15.9%, n=4 animals). nNOS immunochemistry is known to be sensitive to fixation; for example, CA1 PCs are nNOS IHC+ after fixation in 1% PFA, but not 4% PFA^53^. Therefore, we also examined immunolabeling of LINCs after brief 1% PFA, and found a high proportion of LINCS with nNOS immunoreactive somata (349/450, 77.0 ± 9.5%, n=3 animals). **b**) A representative example of somatic colocalization of eYFP with nNOS IHC (closed arrow). Note that not all nNOS IHC+ neurons are labeled with the virus (open arrow), which may be due to the serotype (DJ) of the viral vector used; the lack of viral-labeling of other nNOS IHC+ interneurons, including neurogliaform cells^16,18^, was not due to the Cre or Flpe line used: we crossed cNOS-fDLX mice with a RCE:dual-reporter line, and observed cell populations in areas and numbers consistent with previous literature^15^ (data not shown). **c**) In addition to somatic expression, nNOS can be found in dendrites, including PC dendrites after 1% PFA fixation^53^. In tissue with 1% PFA fixation, we noted instances in which LINC processes were nNOS IHC+ (black arrow) despite having immuno-negative somata (white arrow), indicating that the ~80% of eYFP+ cells with detected somatic nNOS immunoreactivity is a lower-bound of nNOS expression, which underestimates the expression of nNOS in LINCs. Mean±SD. Scale bars=20μm (**b,c**).

**Extended Data Figure 2.**
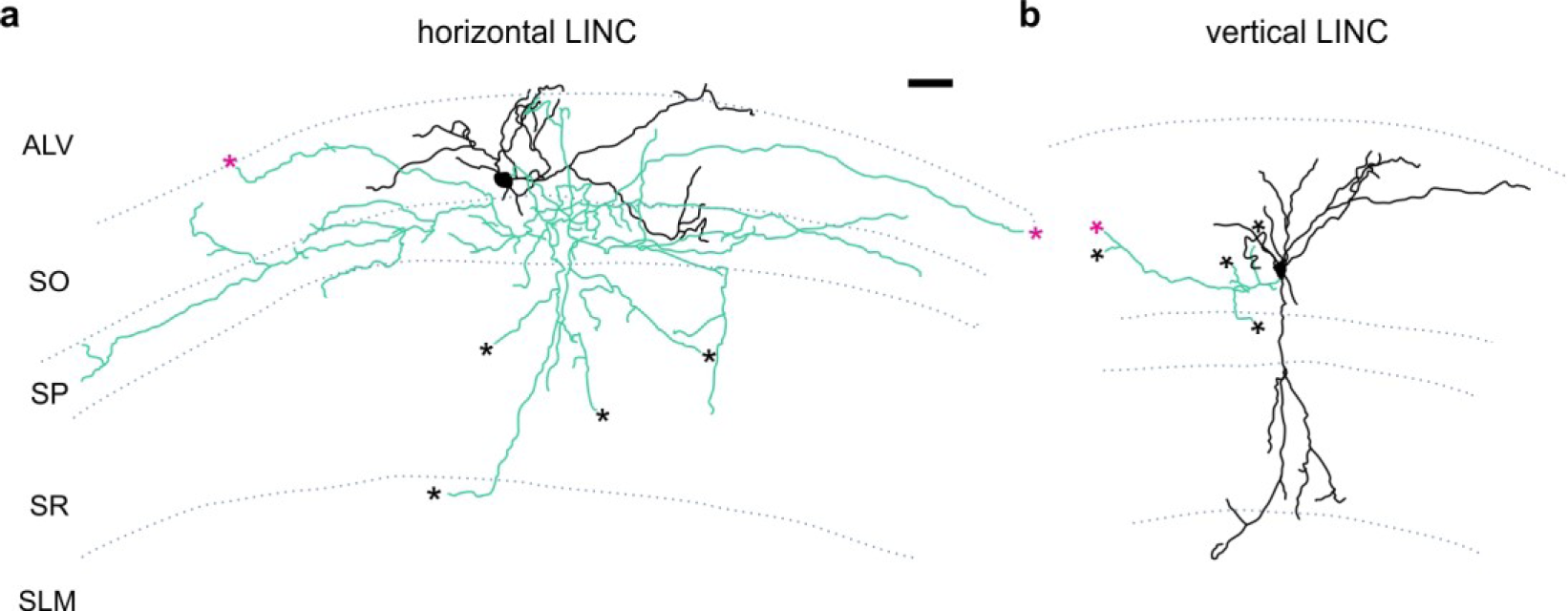
Additional examples of LINC morphology. LINCs filled and recovered in sagittal sections. LINCs show either largely horizontal (**a**) or vertical (**b**) dendrites (black). Note that in addition to local axons (green), filled LINCs show a severed axon (asterisks), suggestive of long-range projections and, unlike previously described hippocampal-septal and back-projection cells, do not show drumstick-like appendages^23,24^. Alveus (AL), stratum oriens (SO), stratum pyramidale (SP), stratum radiatum (SR), stratum lacunosum moleculare (SLM). Scale bar: 50μm (**a,b**).

**Extended Data Figure 3.**
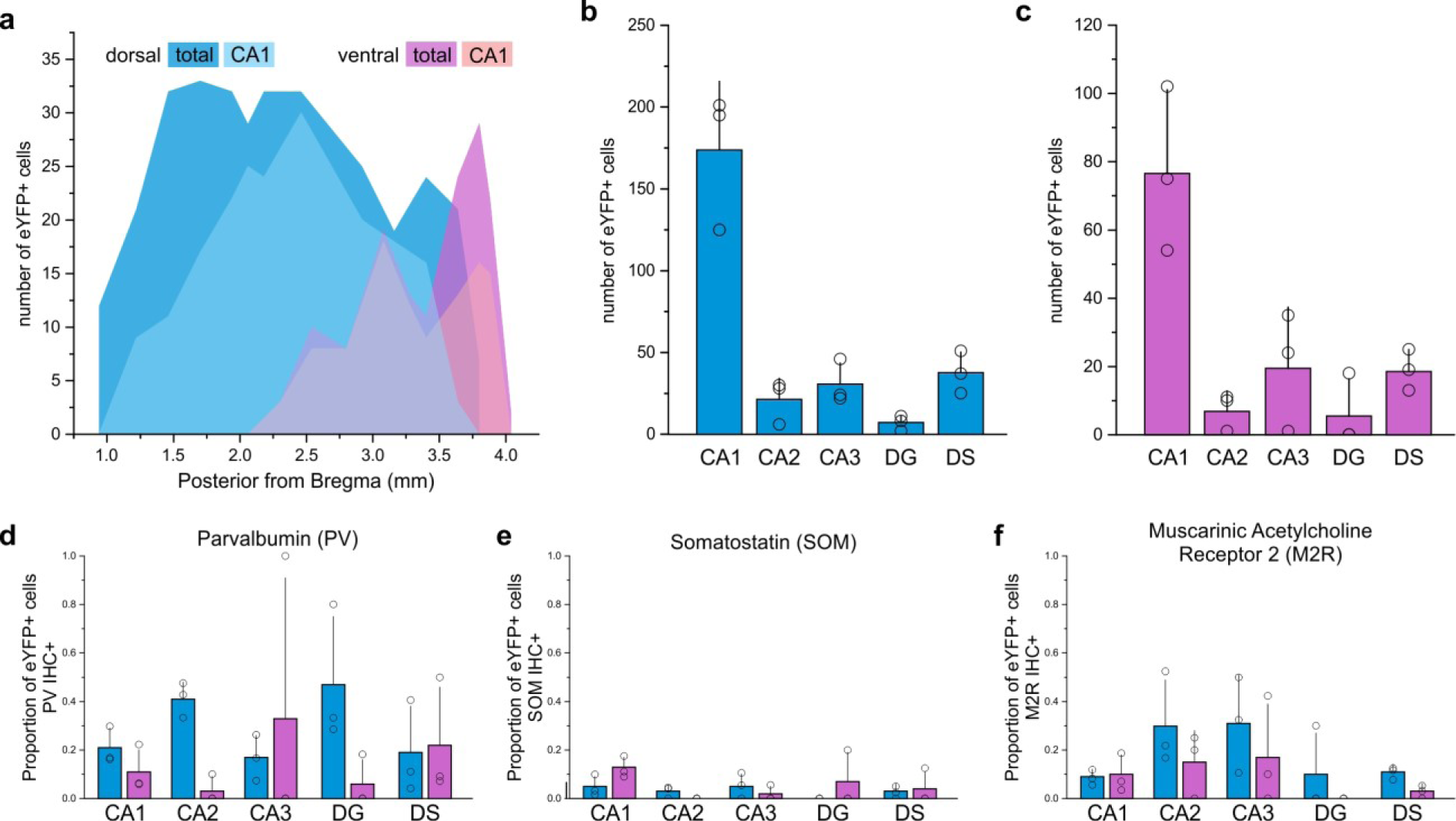
LINCs are found in anterior, intermediate, and posterior regions of the hippocampus. **a)** Distribution of eYFP-labeled cells in an animal after dorsal (blue) and an animal after a more ventral (violet) injection, found in CA1 (light) or all hippocampal subfields (dark). **b-f**) Summary data based on 3 dorsally injected and 3 ventrally injected animals. The majority of eYFP+ cells were found in CA1 (**b**,**c**), but some eYFP+ cells were located in each region of the hippocampal formation following either dorsal (**b**) or ventral (**c**) viral injection. The proportion of eYFP+ cells that colocalize with PV (**d**), SOM (**e**), or M2R (**f**) immunohistochemistry (IHC) following a dorsal (blue) or ventral (violet) viral injection, by hippocampal formation region. Note that the CA1 data (**d-f**) is displayed in the main text, but has also been included here for easy reference. Note also the relatively high proportion of eYFP-labeled cells in the DG that colocalize with PV-immunofluorescence after dorsal virus injection; these cells may correspond to previously noted PV+ nNOS+ cells of the DG, of which little is known^18^. DG: dentate gyrus. DS: dorsal subiculum. Data in **b-f** displayed as mean+SD.

**Extended Data Figure 4.**
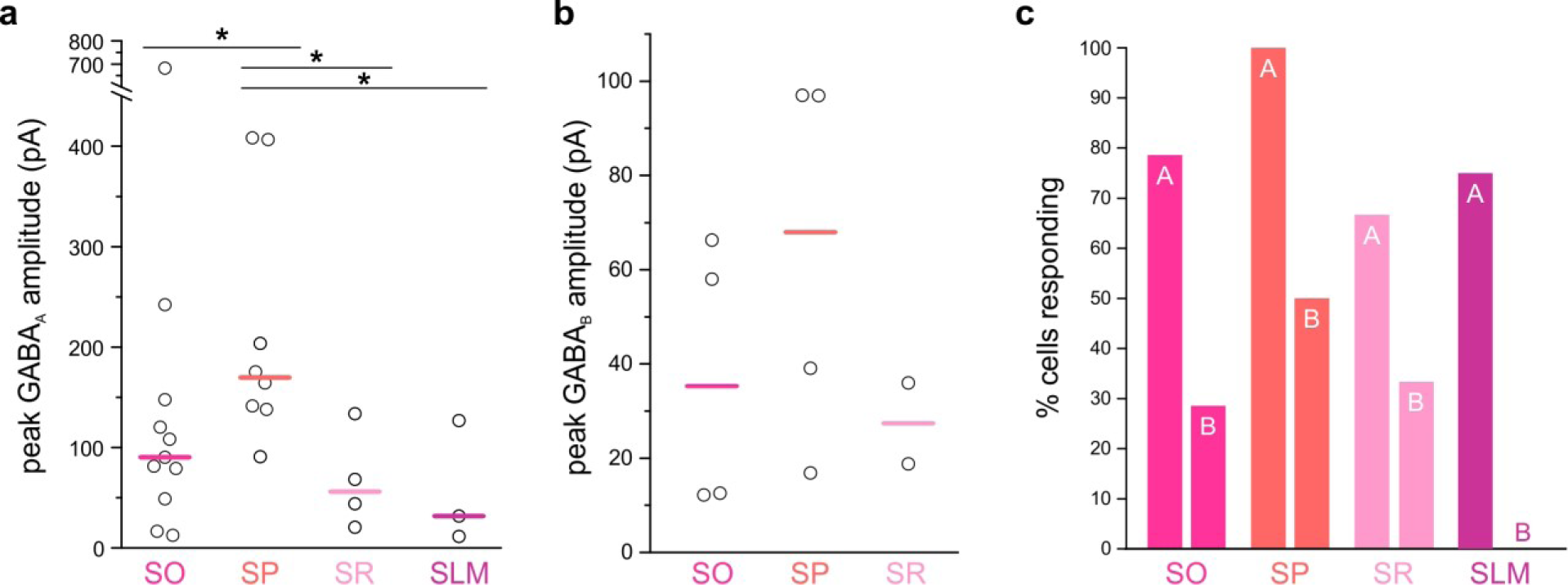
LINCs produce fast and long-lasting inhibitory postsynaptic currents in a variety of inhibitory neurons in CA1. Brief optogenetic activation of LINCs produces postsynaptic inhibitory responses in inhibitory neurons (INs) in CA1 of the hippocampus. **a)** Light-evoked IPSCs were recorded from INs in all strata, with particularly strong GABA_A_-mediated IPSCs recorded from cell bodies in the pyramidal cell layer (SP), (excluding non-responses: Krusall-Wallis (K-W) *p*=0.02; post hoc M-W tests *p*=0.04 (SO v. SP), *p*=0.01 (SP v. SR), *p*=0.03 (SP v. SLM); including non-responses: K-W *p*=0.01; post hoc M-W tests *p*=0.01 (SO v. SP), *p*=0.004 (SP v. SR), *p*=0.01 (SP v. SLM). **b)** GABA_B_ –mediated responses were also evident in many INs (with no statistically significant differences between layers; excluding non-responses: K-W *p*=0.44; including non-responses: K-W *p*=0.26). **c)** The percentage of inhibitory neurons with cell bodies in each layer of CA1 that showed a postsynaptic GABA_A_ (denoted with ‘A’) or GABA_B_ response (‘B’). Notably, every recorded SP interneuron showed a GABA_A_ response (8/8 cells). Note that only cells showing a response are illustrated in panels **a & b**.

**Extended Data Figure 5.**
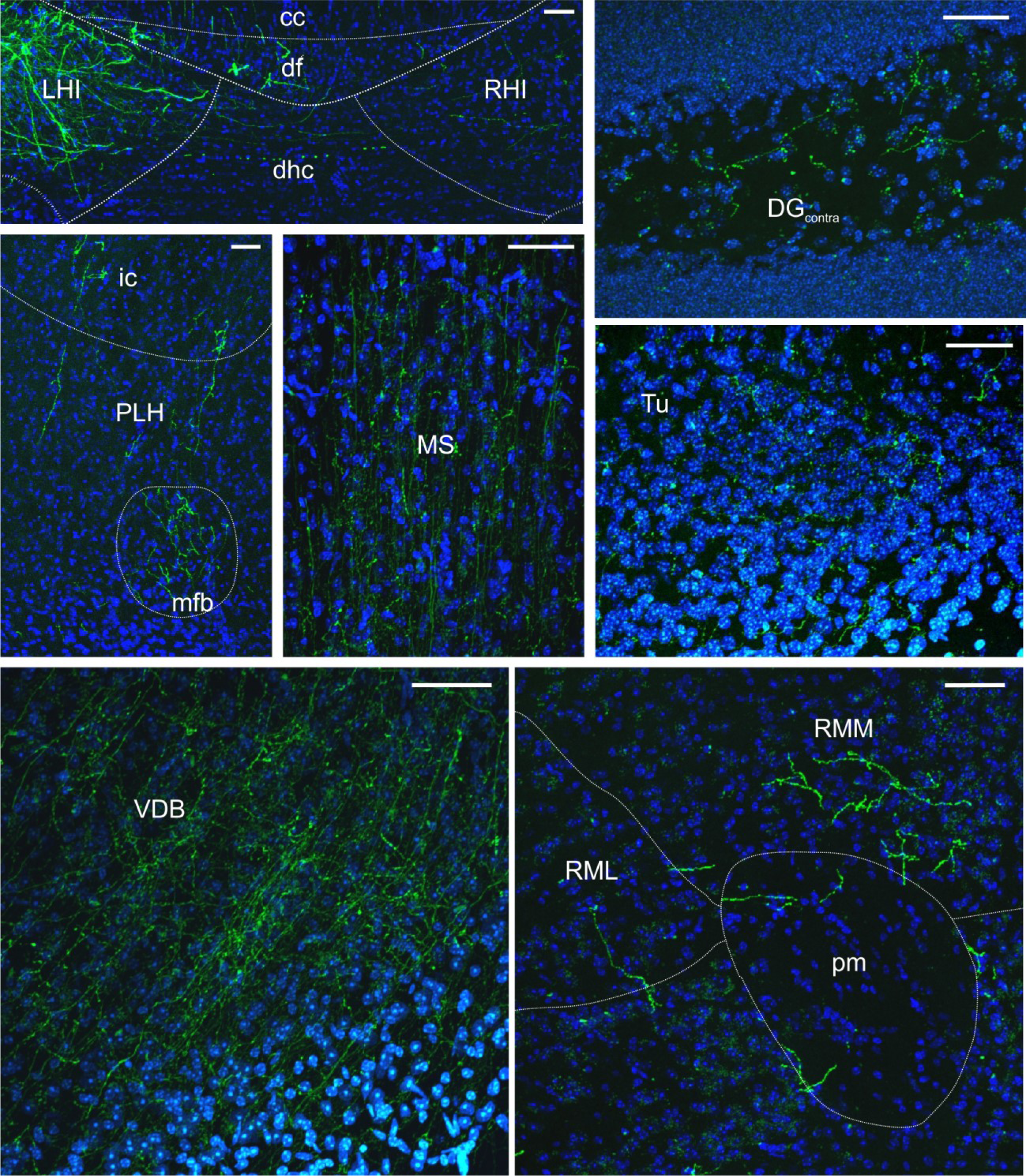
LINCs have long-range projections. Representative images of eYFP+ fibers in various projection regions taken from 50μm coronal sections; neuroanatomical abbreviations and outlines estimated following Franklin & Paxinos mouse brain atlas^72^. DAPI in blue. Corpus callosum (cc), dentate gyrus contralateral (DG_contra_), dorsal fornix (df), dorsal hippocampal commissure (dhc), internal capsule (ic), left hippocampus (LHI), medial forebrain bundle (mfb), medial septum (MS), peduncular part of the lateral hypothalamus (PLH), principal mammillary tract (pm), right hippocampus (RHI), retromammillary nucleus, lateral (RML), retromammillary nucleus, medial (RMM), olfactory tubercle (Tu), vertical limb of the diagonal band of Broca (VDB). Scale bars=50μm.

**Extended Data Figure 6.**
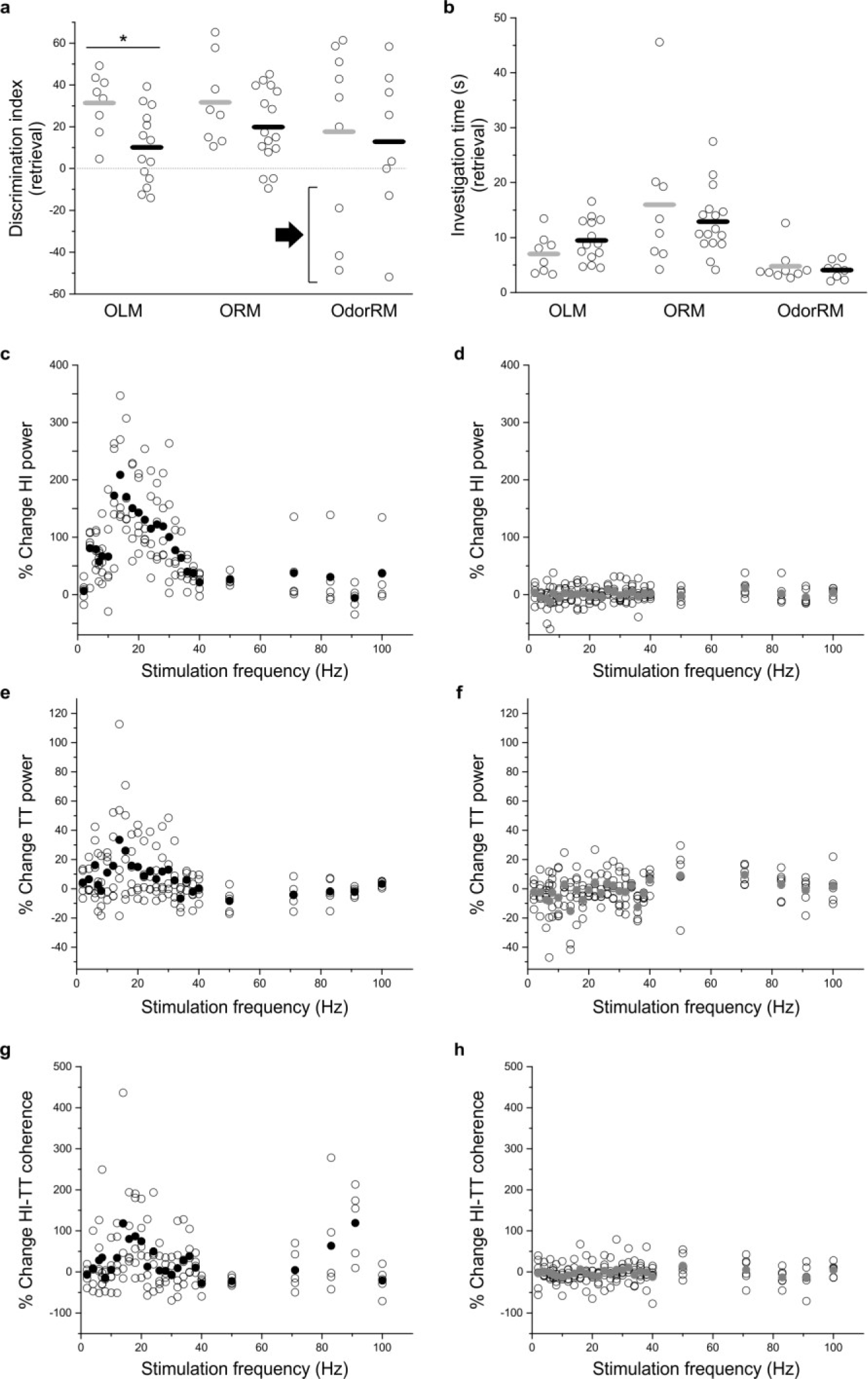
Expansion of *in vivo* behavior and electrophysiology, including individual data points. **a)** Performance (as measured by discrimination index (DI) during retrieval) on the object location, object recognition, and odor recognition memory tasks (OLM, ORM, OdorRM, respectively) in individual opsin+ (black) and opsin-negative (gray) animals (mean: horizontal bar). OLM and ORM data are presented in Fig.5, and are expanded here to display individual animals’ data points, and for easy comparison with the OdorRM test. In the OdorRM test, there was considerable variability in retrieval DI, in both opsin+ and opsin-negative mice, and only modest average DIs (opsin+: 12.8 ± 12.6; opsin-negative: 17.7 ± 14.4; n=8 and n=9 animals, respectively, mean±SEM; no significant difference between genotypes: *p*=0.67, M-W). This appeared to potentially be due to inter-mouse variability regarding which odor was preferred (i.e., novel^62^ vs familiar^73^; note the large negative DIs for some animals, arrow). When the sign (positive/negative) of the DI was not taken into account, OdorRM average DIs were markedly higher in both genotypes (opsin+: 29.0 ± 7.8, opsin-15 negative: 41.9 ± 5.1, not illustrated), but there remained no significant difference between genotypes (opsin+ vs. opsin-negative, *p*=0.27, M-W). **b)** Time spent investigating the objects during retrieval. Animals that investigated fewer than 3s (OLM, ORM) or 2s (OdorRM) were excluded from analysis and are not shown. **c-h**) Percent change in hippocampal (**c,d**) or tenia tecta (**e,f**) power or hippocampal-tenia tecta coherence (**g,h**) when LINCs were stimulated at a variety of frequencies in individual opsin+ (**c,e,g**) and opsin-negative (**d,f,g**) animals. Mean: closed circles (**c-h**). Note y-axis changes for the different plots, and that this figure expands upon Fig.5, to show individual animals’ data points.

**Extended Data Figure 7.**
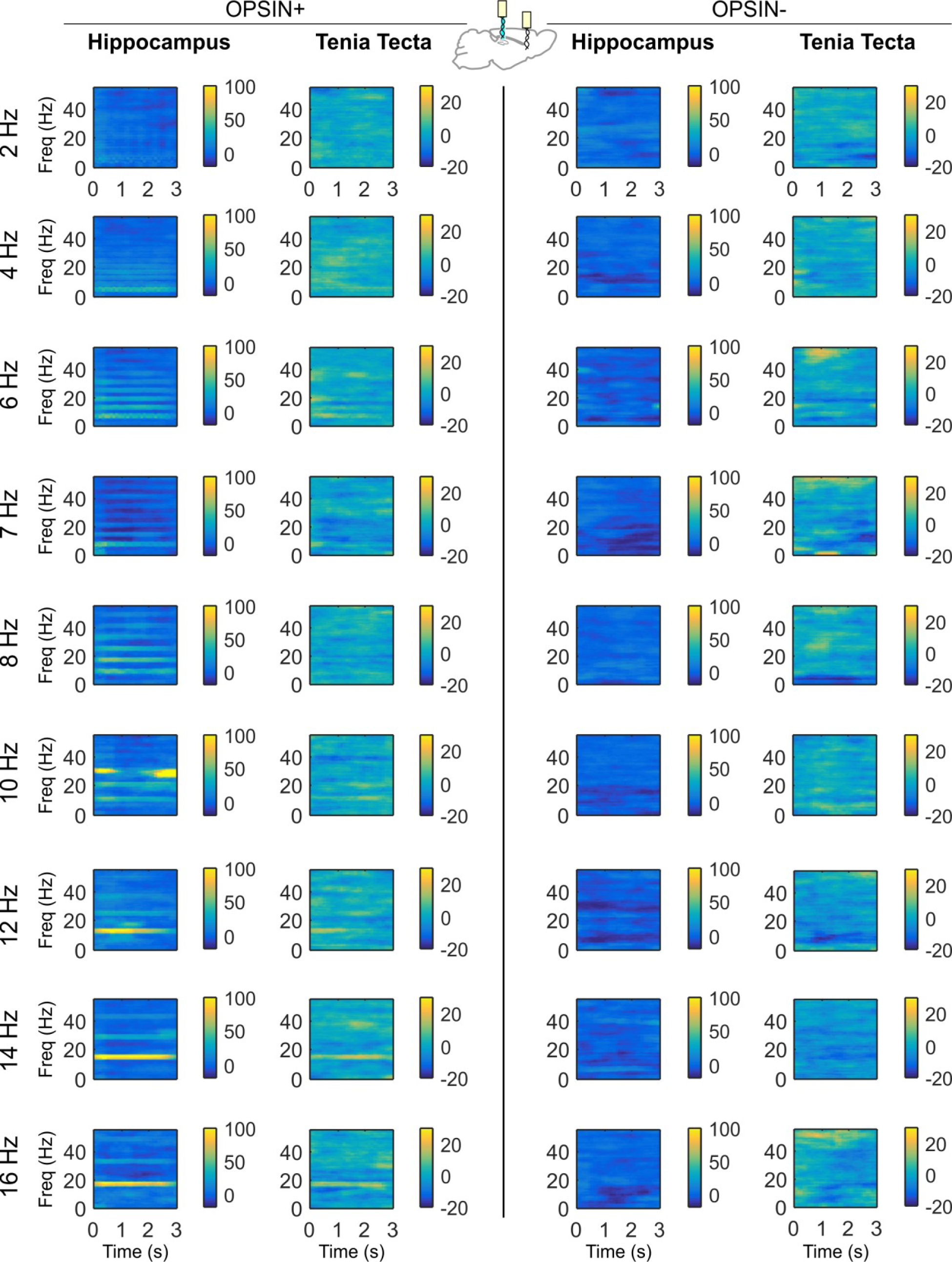
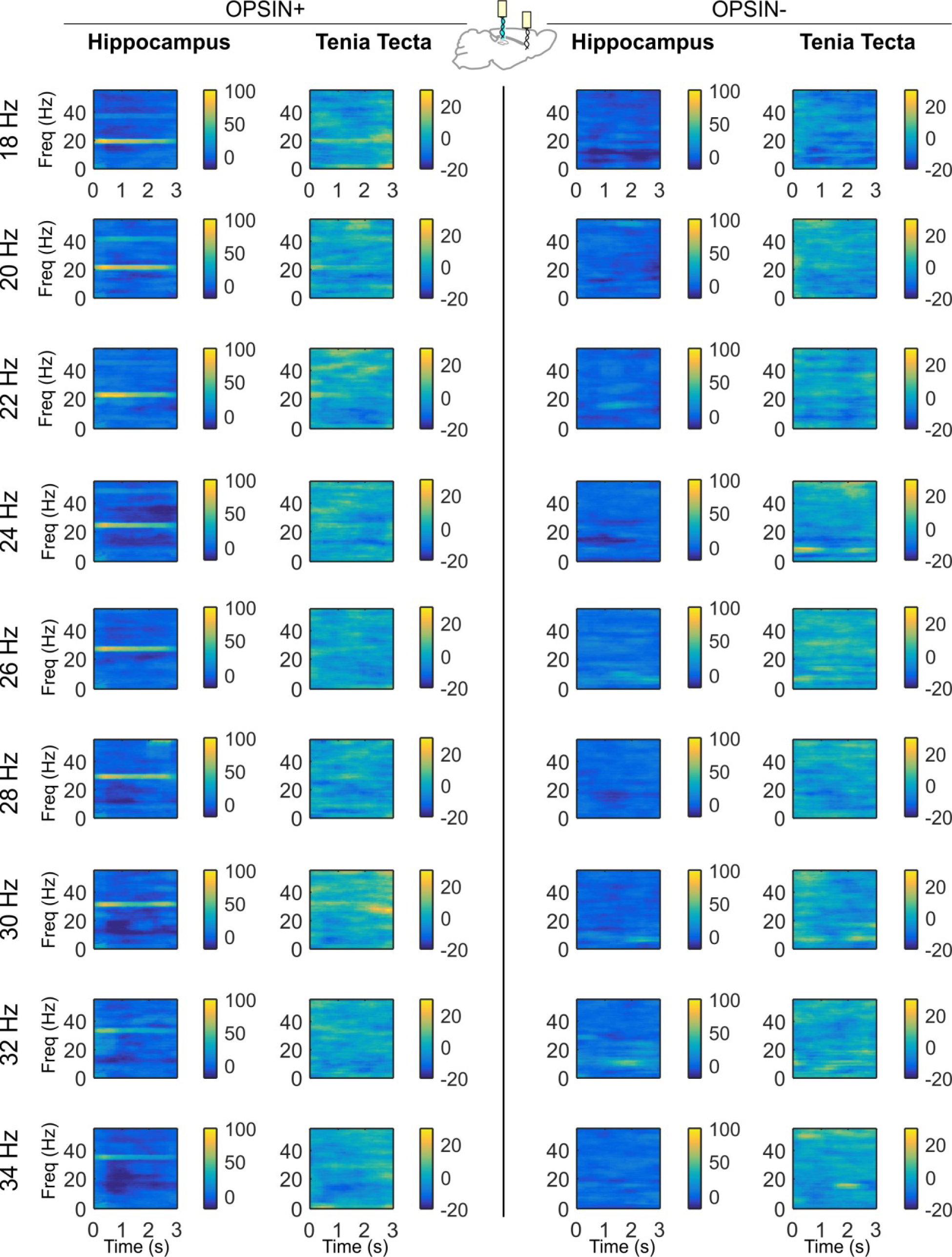
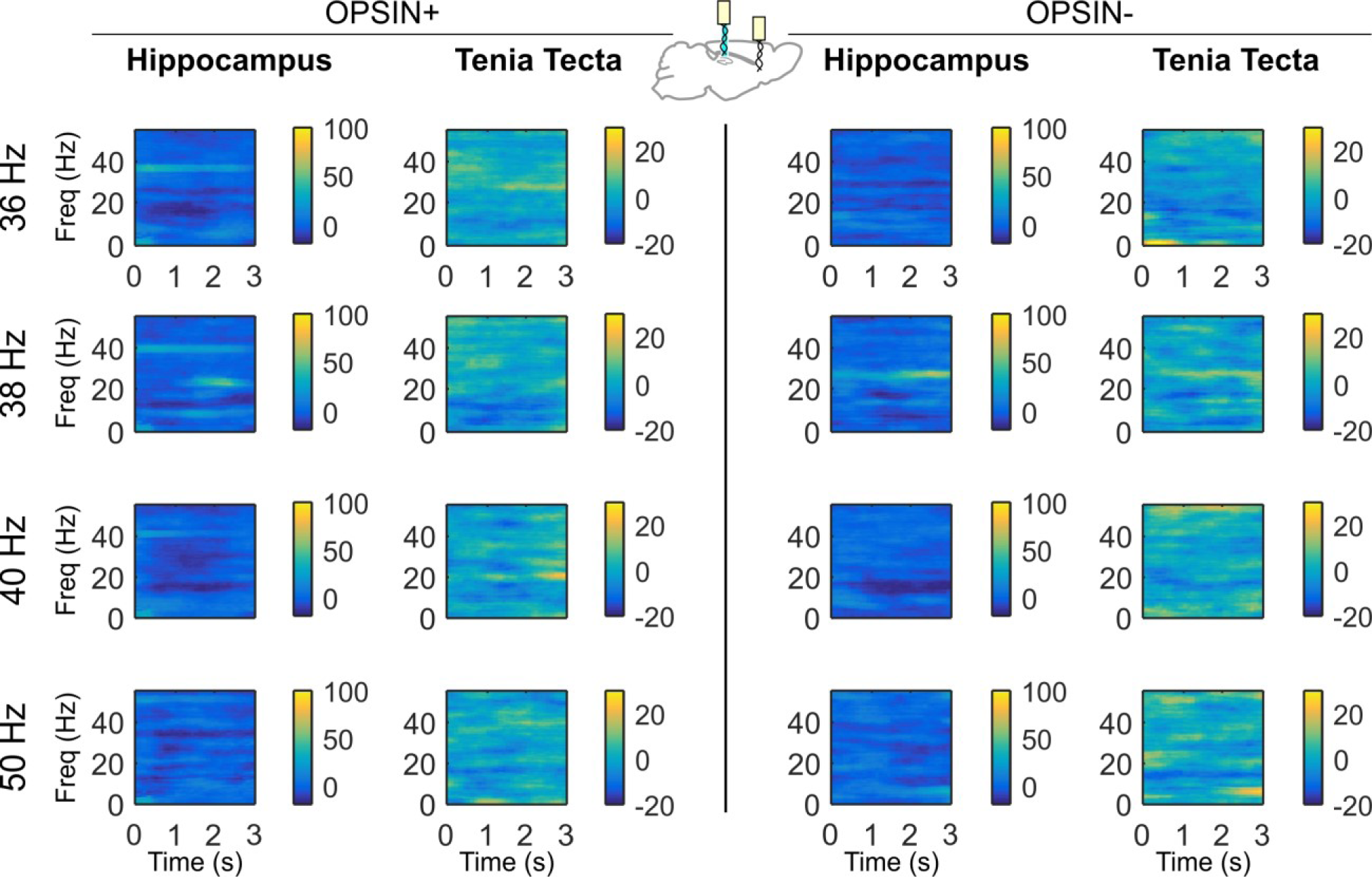
Differential spectrograms showing increases in hippocampal and TT power with different stimulation frequencies. Percent change in power at the stimulation frequency shown in opsin+ (left, average of n=5 animals) and opsin-negative (right, average of n=6) animals in the hippocampus and tenia tecta during light delivery. Light delivered to the hippocampus between 4Hz and 40Hz produced strong increases in power in the hippocampus at the stimulation frequency. Light delivered to the hippocampus between 12Hz and 20Hz also produced noticeable increases in power at that stimulation frequency in the tenia tecta.

## Acknowledgements

The authors would like to thank Gord Fishell and his lab, including Robert Machold, Goichi Miyoshi, and Rowena Turnbull for providing the Dlx5/6-Flpe and RCE:DUAL mouse lines, and Karl Deisseroth and Charu Ramakrishnan for the intersectional virus. We thank the University of Minnesota University Imaging Centers for tissue clearing and imaging and technical support, with special assistance from Director Mark Sanders. Additional thanks to the entire Krook-Magnuson lab, in particular, to Casey Xamonthiene for expertise in tissue processing, Zachary Zeidler and Zachary Montes for assistance with behavioral experiments, Chris Krook-Magnuson for MATLAB programming assistance, and Caara H. Leintz for early assistance with stereotactic surgeries. This work was supported by NIH grants R01-NS104071 (to EKM) and F31-NS105457 (to ZCW), the University of Minnesota’s MnDRIVE (Minnesota’s Discovery, Research and Innovation Economy) initiative (to EKM & ZCW), and a McKnight Land-Grant Professorship (to EKM).

## Author Contributions

Study concept and design: EKM, ZCW. Acquisition of data: ZCW, MRT. Analysis and interpretation of data: ZCW, MRT, EKM. Statistical analysis: ZCW, EKM. Drafting of the manuscript: ZCW, MRT, EKM. Study supervision: EKM.

